# The molecular recognition of phosphatidic acid by an amphipathic helix in Opi1

**DOI:** 10.1101/250019

**Authors:** Harald F. Hofbauer, Michael Gecht, Sabine C. Fischer, Anja Seybert, Achilleas S. Frangakis, Ernst H. K. Stelzer, Roberto Covino, Gerhard Hummer, Robert Ernst

## Abstract

A key event in cellular physiology is the decision between membrane biogenesis and fat storage. Phosphatidic acid (PA) is an important lipid intermediate and signaling lipid at the branch point of these pathways and constantly monitored by the transcriptional repressor Opi1 to orchestrate lipid metabolism. Here, we report on the mechanism of membrane recognition by Opi1 and identify an amphipathic helix (AH) for the selective binding to membranes containing PA over phosphatidylserine (PS). The insertion of the AH into the hydrophobic core of the membrane renders Opi1 sensitive to the lipid acyl chain composition as an important factor contributing to the regulation of membrane biogenesis. Based on these findings, we rationally designed the membrane binding properties of Opi1 to control its responsiveness in the physiological context. Using extensive molecular dynamics (MD) simulations, we identified two PA-selective three-finger grips that tightly bind the phosphate headgroup, while interacting less intimately and more transiently with PS. This work establishes lipid headgroup selectivity as a new feature in the family of AH-containing membrane property sensors.

## Introduction (concise, not limited, no subheadings)

Lipids are actively involved in cellular signaling and serve as major determinants of the organellar identity (Holthuis & Menon, 2014; Bigay & Antonny, 2012). Numerous molecular processes occur at the surfaces of organelles, and the selective recruitment of cytosolic effectors to specific target membranes is crucial to control lipid metabolism, vesicular transport, and cellular signaling (Antonny, 2011; Jacquemyn *et al*, 2017; Odorizzi *et al*, 2000). The organelles of eukaryotic cells are composed of hundreds of lipid species (Zinser *et al*, 1991; Klemm *et al*, 2009; Gerl *et al*, 2012; Ejsing *et al*, 2009).Despite a continuous exchange of membrane material (Holthuis & Levine, 2005) organelles maintain their characteristic lipid compositions and surface properties (Antonny *et al*, 2015; Bigay & Antonny, 2012; De Kroon *et al*, 2013). A particularly powerful mechanism of membrane homeostasis is a feedback control by membrane-associated transcription factors and transcriptional programs that can either sense the level of individual lipids such as cholesterol (Goldstein *et al*, 2006) or phosphoinositides (PIPs) (Laplante & Sabatini, 2012), or even respond to bulk physicochemical membrane properties (Covino *et al*, 2016; Halbleib *et al*, 2017).

The decision to direct lipid precursors to either membrane biogenesis or fat storage represents a key regulatory step in cellular physiology. The transcriptional programs underlying these processes must be carefully coordinated and tightly controlled (Puth *et al*, 2015; Henry *et al*, 2012). Phosphatidic acid (PA) is a class of glycerophospholipids at the branch point of membrane lipid biosynthesis and triacylglycerol (TAG) production (Ernst *et al*, 2016; Athenstaedt & Daum, 1999). As hydrolysis products of phospholipase D1 and D2, PA lipids act as second messengers (Shin & Loewen, 2011). This signaling function is conserved in yeast (Loewen, 2004), plants (Testerink & Munnik, 2011), and mammals (Laplante & Sabatini, 2012; Wang *et al*, 2006). Moreover, PA lipids have also been implicated in the regulation of phosphatidylcholine (PC) biosynthesis by modulating the membrane recruitment of CCTa (PCYT1A in humans) (Cornell, 2016) and in the fission of mitochondria via Drp1 (Adachi *et al*, 2016). A misregulated metabolism of PA has been firmly implicated in cancer biology (Foster, 2009; Laplante & Sabatini, 2012), but the molecular mechanisms of PA recognition remain elusive (Liu *et al*, 2013).

Given the central position of PA lipids in cellular physiology and the lipid metabolic network (Henry *et al*, 2012), it is not surprising that cells established mechanisms to monitor the level of PA. Opi1 is a soluble, transcriptional repressor in *S. cerevisiae* controlling the expression of lipid biosynthetic genes containing an upstream activating sequence responsive to inositol (UASINO). These genes are involved in the production of the major glycerophospholipid classes PC, phosphatidylethanolamine (PE), phosphatidylinositol (PI), and phosphatidylserine (PS), for which PA lipids serve as precursors (Henry *et al*, 2012). When the level of PA is high, Opi1 directly binds to PA at the ER membrane and is prevented from entering the nucleus, thereby allowing for the expression of UASINO target genes (Fig 1A) (Loewen, 2004). When PA is consumed, Opi1 is released from the ER, translocates into the nucleus due to its nuclear localization sequence (KRQK, aa 109-112), and represses membrane biogenesis genes (Loewen, 2004) (Fig 1B). The direct binding of Opi1 to PA-rich membranes is assisted by the tail-anchored VAP ortholog protein Scs2 that binds the Opi1 FFAT domain (two phenylalanines in an acidic tract; aa 193-204) and acts as a co-receptor in the ER membrane (Fig 1A,C) (Loewen *et al*, 2003; Loewen, 2004). Inositol is a master regulator of this pathway (Jesch *et al*, 2005): When inositol is present in the medium, PA lipids are converted to PI and Opi1 represses its target genes, including *INO1*, which encodes the rate-limiting enzyme for inositol biosynthesis (Graves & Henry, 2000) (Fig 1B). Loss of Scs2, the co-receptor for Opi1, causes a constitutive repression of *INO1* and thus inositol auxotrophy (Loewen, 2004; Gaspar *et al*, 2017). Loss of Opi1, on the contrary, results in the de-repression of lipid biosynthetic genes as evidenced by the overproduction and secretion of inositol from cells, the so called Opi^−^ phenotype (Greenberg et al, 1982). The expression level of *OPI1* underlies a self-regulatory feedback loop, as its promoter also contains a UASINO sequence (Schüller *et al*, 1995).

**Figure 1:**
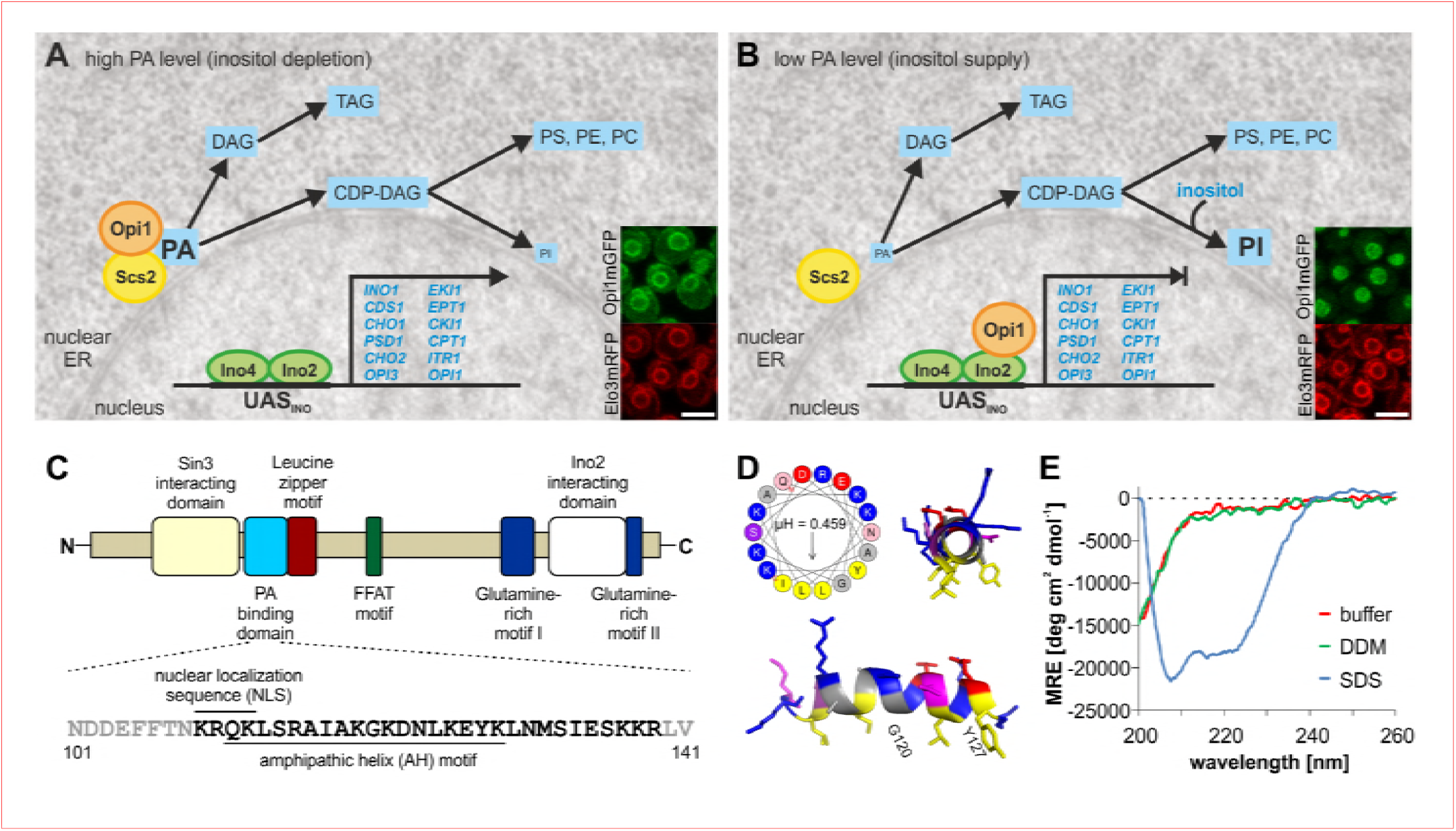
Opi1 uses an amphipathic helix (AH) to control glycerophospholipid metabolism. A,B The regulatory role of Opi1. When the level of PA is high in the nuclear ER, Opi1 is bound to the ER thereby facilitating membrane biogenesis (A). When the level of PA is low, Opi1 localizes to the nucleus and represses its target genes (B). Scs2 binds the Scs2 interacting FFAT motif in Opi1 to assist membrane recruitment. Inlets show live cell microscopic images of Opi1-mGFP localization with Elo3-mRFP as a marker for the ER. Scale bar = 5 μm. C Domain organization of Opi1 (404 aa) highlights the PA interacting motif (aa 109-138; light blue), a leucine zipper motif (aa 139-160, brown), the Scs2 interacting FFAT motif representing an acidic tract with two phenylalanines (aa 193-204, green). The expanded view shows the amino acid sequence of the nuclear localization sequence (aa109-112) and the PA binding domain forming a putative AH (aa 111-128). D Visualization of the AH using the Heliquest tool (Gautier et al., 2008) and the PyMOL^TM^ Molecular Graphics System for front and side views of the helix matching the color code of the Heliquest tool. E CD spectroscopic analysis of the Opi1^111-128^ synthetic peptide comprising the putative AH recorded in sodium phosphate buffer in the absence of detergent (buffer) or in the presence of either 20 mM dodecyl maltoside (DDM) or 20 mM sodium dodecyl sulfate (SDS).

Recent studies have suggested that Opi1 binding to PA is not only controlled by the level of anionic PA lipids interacting with basic residues in the PA binding domain of Opi1 (Opi1^111-128^), but also by the lipid acyl chain composition (Hofbauer *et al*, 2014; Kassas *et al*, 2017; Putta *et al*, 2016). These observations provided a new perspective on the regulation of membrane biogenesis with Opi1 mediating an intricate crosstalk between fatty acid metabolism and glycerophospholipid biosynthesis. In fact, it was postulated that the PA binding domain of Opi1 might form an amphipathic helix (AH) (Ganesan *et al*, 2015) to engage in interactions both with the lipid headgroups of PA and with the hydrophobic core of the membrane. While the preference of the Opi1 PA binding domain for PA-containing and loosely packed membranes provided supporting evidence (Kassas *et al*, 2017; Putta *et al*, 2016), this possibility was not directly tested.

Given the simplicity of the negatively charged phosphate headgroup of PA, a particularly puzzling question remains: How do proteins distinguish between PA and other anionic lipids such as PS to confer specificity? In contrast to the phosphoesters in PS, the phosphomonoester of PA can be deprotonated twice, yielding ionization states of either -1 or -2 (Kooijman *et al*, 2005). The pKa for the second deprotonation of PA is close to the physiological pH and greatly affected by the membrane context (Kooijman *et al*, 2005). The phosphate moiety of PS can be deprotonated only once, yielding an ionization state of -1 at physiological pH. Based on these findings, it was proposed that the intracellular pH provides an additional cue for the Opi1 regulatory system (Young *et al*, 2010; Shin & Loewen, 2011). Despite these intriguing findings, it is still unclear if specific structural features could endow a protein with an inherent selectivity for PA.

Here we present our efforts to better understand the molecular underpinnings of PA recognition by the Opi1 regulatory system. We experimentally validate the presence of an AH in the basic PA-binding region of Opi1 and establish the tuning of interfacial hydrophobicity as a powerful tool to manipulate membrane binding *in vitro* and membrane-dependent signaling *in vivo*. Using extensive atomistic molecular dynamics (MD), we identify two PA-selective three-finger grips each formed by three basic residues on one side of the AH as a robust mechanism for selective PA binding. Intriguingly, lysine and arginine residues have non-equivalent functions in establishing PA selectivity, thereby excluding that PA recognition is solely dictated by electrostatics. This work establishes headgroup selectivity of AHs as crucial contributors to the regulation of membrane biogenesis in yeast and to membrane recognition processes in general.

## Results

In order to characterize the details of membrane recognition by Opi1, we focused our attention on the PA binding domain (Opi1^111-128^) (Loewen, 2004) adjacent to the nuclear localization sequence (KRQK, Opi1^109-112^) (Fig 1C, Fig EV1A). HeliQuest analysis (Gautier *et al*, 2008) of this region revealed a putative amphipathic helix (AH) with a small hydrophobic and a large hydrophilic face and several basic amino acid residues, which are crucial for PA-binding (Loewen, 2004) (Fig 1D). This arrangement of hydrophobic and basic residues in the predicted AH of Opi1 resembles the AH of the membrane-sensor motif of Spo20 (Spo20^62-79^) (Fig EV1B), but differs significantly from the N-terminal AH of the PA-converting phosphatidate phosphatase Pah1 (Karanasios *et al*, 2010) (Fig EV1C). While the AH of Pah1 (Pah1*1-18*) has a serine-rich hydrophilic face with only two basic residues, the AHs of both Opi1 and Spo20 feature high densities of basic residues (Fig EV1A-C). Using CD spectroscopy, we characterized the secondary structure of a synthetic peptide corresponding to the predicted AH (Opi1^111-128^). The peptide was unstructured in aqueous buffer even in the presence of a hydrophobic matrix provided by dodecylmaltoside (DDM) micelles (Fig 1E). Only in the presence of sodium dodecyl sulfate (SDS) the peptide adopted a helical secondary structure. These observations suggest that the PA-interacting motif of Opi1 has only a weak propensity to form an AH, if at all.

We hypothesized that the AH of Opi1 might form only upon membrane binding, similar to membrane-active antimicrobial peptides (Ladokhin & White, 1999; Shai, 1999) and amphipathic lipid packing sensor (ALPS) motifs (Drin & Antonny, 2010; Antonny, 2011). Because PA supports membrane binding of Opi1 (Loewen, 2004), we studied the secondary structure of the synthetic Opi1^111-128^ peptide in the presence of liposomes with a PC-based matrix and with increasing molar fractions of PA (Fig 2A). The lipid mixtures were prepared from stocks of 1,2-dioleoyl-*sn*-glycero-3-phosphocholine (DOPC), 1-palmitoyl-2-oleoyl-*sn*-glycero-3-phosphocholine (POPC) and 1-palmitoyl-2-oleoyl-*sn*-glycero-3-phosphate (POPA) to generate liposomes differing in the lipid headgroup composition, but not in the lipid acyl chain composition. The resulting liposomes were composed of lipids with 75% monounsaturated and 25% saturated lipid acyl chains to match the molecular packing density of the ER (Kaiser *et al*, 2011). Liposome diameters of 180-200 nm, determined using NanoSight technology, and polydispersity indices below 0.1 suggest a monodisperse preparation of liposomes. Liposomes were inspected by cryo-electron microscopy to further validate the quality of the preparation (Fig EV2D). CD spectroscopy revealed that the Opi1^111-128^ peptide is unstructured in the presence of liposomes composed of only PC lipids, but adopts an alpha-helical conformation in the presence of PA-containing liposomes. The alpha-helical signature of the CD-spectrum increased with the molar fraction of PA in the liposomes (Fig 2A). These data suggested that the PA binding domain of Opi1 is unstructured in solution but folds into an AH upon PA-dependent membrane binding.

**Figure 2:**
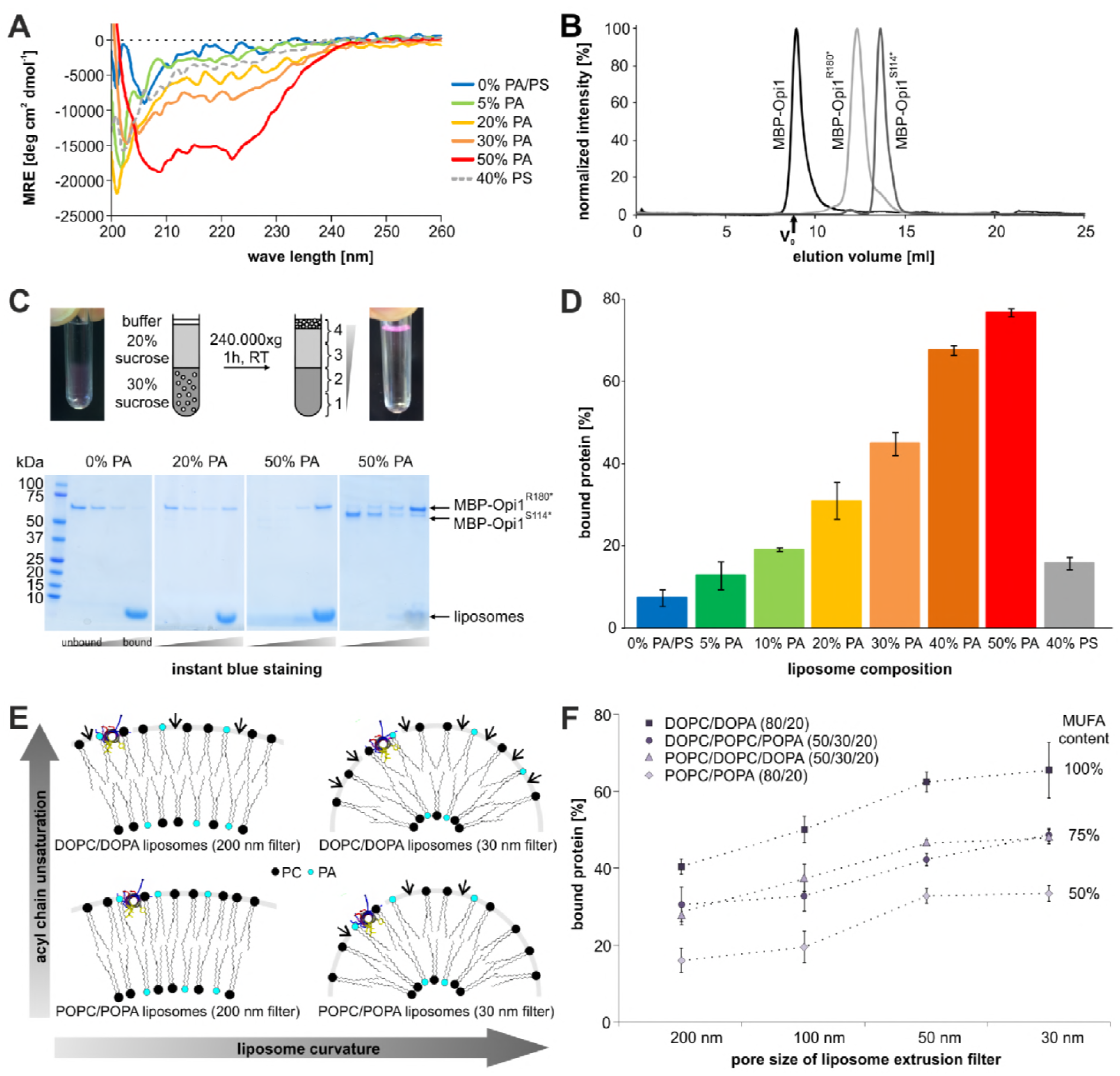
The AH of Opi1 senses PA, lipid packing, and membrane curvature. A The Opi1^111-128^ synthetic peptide folds upon binding to PA-containing membranes as revealed by CD spectroscopic analysis. The peptide comprising the putative AH is unstructured in the presence of liposomes composed only of PC (0% PA/PS), but the helical content increases with the proportion of PA in the liposomes. As control, CD spectra were recorded with liposomes containing 40 mol% PS (dashed grey line). B The indicated MBP-Opi1 fusion constructs were purified by affinity chromatography and subsequently analyzed by size exclusion chromatography. The void volume (V_0_) of the Superdex 200 Increase 10/300 GL column is 8.9 ml. C Indicated MBP-Opi1 variants were incubated with liposomes containing different PA concentrations. After centrifugation in a discontinuous sucrose gradient to float all liposomes (as schematically illustrated) the gradient was fractionated (as indicated). The fractions were subjected to SDS-PAGE to quantify the percentage of Opi1 found in the top fraction of the gradient. D Quantification of liposome flotation assays. The percentage of Opi1 found in the top fraction of the gradient was determined by densiometry. The bar diagrams show the average of 2 independent experiments for 10%, 40% and 50% PA, 3 independent experiments for 5% and 30% PA, 9 independent experiments for 0% PA/PS and 40% PS and 11 independent experiments for 20% PA. Error bars represent the standard deviation. E Schematic illustration of the impact of membrane curvature and the unsaturation degree of lipid acyl chains on the frequency of interfacial voids. Arrows indicate potential membrane penetration sites for Opi1’s AH. F The binding of MBP-Opi1^180*^ (0.63 μM) to different liposomal preparations (2.1 mM lipids) was analyzed by liposome floatation assays as in Figure 2D. The proportion of unsaturated lipid acyl chains was varied by using the indicated species of PA and PC, while the radii of the liposomes were modified by extrusion through polycarbonate filters with indicated pore sizes. Dashed lines are included in the graph to underscore the role of membrane curvature. Each data point represents the average of at least 3 independent experiments with the error bars representing the standard deviation.

Previous work on the membrane-sensor motif of Spo20 had suggested that it contains a PA-selective membrane-binding region (Spo20^62-79^) (Nakanishi, 2004; Kassas *et al*, 2012). GFP fusion proteins based on Spo2051-91 have been established (Nakanishi, 2004) and optimized (Kassas *et al*, 2012; Zhang et al, 2014) to act as biosensors for PA in living cells. More recent studies by Antonny and coworkers reported that Spo20 does not efficiently distinguish between the anionic lipids, PA, PS and PIPs (Horchani *et al*, 2014). Instead, they found that the membrane binding of Spo2051-91 relies on the negative charge density on the surface of the liposomes. The net charge of the PA headgroup can be -1 or -2, while the net charge of PS is only -1. When compensating for this charge difference by increasing the PS concentration, Spo2051-91 did no longer show any preference for PA-containing liposomes (Horchani *et al*, 2014). Given the overall similarity of the AH regions of Spo20 and Opi1 (Fig EV1A,B), we performed a similar control experiment with Opi1^111-128^. CD spectroscopy revealed that the helical content of the AH peptide was higher in the presence of liposomes with 20 mol% PA than in the presence of liposomes with 40 mol% PS (Fig 2A). These results suggest that the AH of Opi1 has an inherent selectivity to PA, which is missing in the AH of Spo20 despite the overall similarity.

Having shown that the isolated PA binding region of Opi1 (Opi1^111-128^) folds upon binding to PA-containing membranes, we aimed at determining the membrane recognition process also in the context of the Opi1 protein. To this end, we generated fusion constructs of MBP and Opi1 (full length and C-terminal truncations) for the heterologous overproduction in *Escherichia coli*. After affinity-purification of MBP-Opi1 (Loewen, 2004; Sreenivas & Carman, 2003) (Fig EV2A), we analyzed the oligomeric state of the fusion proteins by size exclusion chromatography. Full-length MBP-Opi1 eluted as higher oligomer in the void volume of a Superdex 200 column (Fig 2B). The truncation variant MBP-Opi1R^180*^ (predicted molecular weight: 63.3 kDa) containing both the PA-binding region and the functionally important leucine zipper motif (Opi1^139-160^) (Fig 1C) (White et al, 1991) eluted as oligomer, while the MBP-Opi1^S114*^ variant (55.9 kDa) lacking these two domains eluted as a monomeric protein (Fig 2B). The elution fractions of affinity-purified MBP-Opi1R^180*^ were analyzed by SDS-PAGE and only the peak fraction from the size exclusion chromatography column (Fig EV2A-C) was used for subsequent binding assays.

The membrane binding and lipid selectivity of Opi1 was investigated using liposome flotation assays (Fig 2C). MBP-Opi1R^180*^ was incubated in the presence of PC-based liposomes containing different molar fractions of PA or PS but exhibiting identical lipid acyl chain compositions. After flotation of the liposomes in a discontinuous sucrose gradient, four equal fractions were retrieved from the gradient and subjected to SDS-PAGE. The percentage of membrane-bound Opi1 in the top fraction was quantified by densiometry using Instant Blue^TM^, which stains proteins and lipids, thus providing a convenient way to validate the successful flotation of the liposomes (Fig 2C). Consistent with our findings on the isolated PA-binding region (Opi1^111-128^) by CD spectroscopy (Fig 2A) we found that the liposome binding of MBP-Opi1R^180*^ increases with the molar fraction of PA in the lipid bilayer (Fig 2C,D). Membrane binding was fully dependent on the PA binding region and the leucine zipper, as the MBP-Opi1^S114*^ fusion protein failed to bind to PA-containing liposomes (Fig 2C, right panel). The striking selectivity of Opi1 for PA- over PS-containing membranes was underscored by the observation that liposomes with 20 mol% PA bound Opi1 more efficiently than liposomes with 40 mol% PS (Fig 2D). Thus, even when compensating for the maximum possible difference of the headgroup charge per area of membrane, Opi1 favors binding to PA- over PS-containing membranes, strongly suggesting that it endows specific features for PA selectivity.

If Opi1 recognizes PA-containing membranes by folding an AH into the lipid bilayer, the membrane binding should be affected by the membrane curvature and the molecular lipid packing density of the lipids. Interfacial membrane voids, also referred to as lipid packing defects, caused by membrane curvature and loosely packing lipids should facilitate the insertion of hydrophobic side chains into the core of the lipid bilayer and thus the folding of the AH (Drin & Antonny, 2010; Antonny, 2011). Consistent with this hypothesis, we find that membrane binding of MBP-Opi1R^180*^ increased with the membrane curvature and the molar fraction of monounsaturated acyl chains in the bilayer (Fig 2E). MBP-Opi1R^180*^ was incubated with liposomes containing 20 mol% PA but differing either in their membrane curvature or their acyl chain composition. The curvature of the liposomes was adjusted by stepwise extrusions through polycarbonate filters with decreasing pore sizes from 200 nm to 30 nm. The lipid compositions were chosen to yield different proportions of esterified monounsaturated fatty acids (MUFA content) between 50% and 100%. After floating the liposomes in a sucrose step gradient, the fraction of MBP-Opi1^180*^ co-floating with the liposomes was determined. The binding of MBP-Opi1^180*^ to liposomes was favored both by an increased membrane curvature and by an increased proportion of monounsaturated lipid acyl chains in the lipid bilayer (Fig 2F). Thus, even though the AH of Opi1 differs from classical ALPS motifs (Drin & Antonny, 2010) by the presence of basic residues in the hydrophilic face, it shares their sensitivity to membrane curvature and the lipid acyl chain composition. However, whereas the overall proportion of saturated and unsaturated lipid acyl chains had a significant impact on the membrane recruitment of MBP-Opi1^180*^ (Fig 2F) the precise position of the saturated lipid acyl chains appeared irrelevant. When 20 mol% of POPA were substituted by 20 mol% 1,2-dioleoyl-*sn*-glycero-3-phosphate (DOPA) and the respective changes of the lipid acyl chain composition was compensated by changes in the POPC and DOPC content to maintain the overall proportion of unsaturated lipid acyl chains at 75 mol%, the binding of MBP-Opi1^180*^ was barely affected, if at all (Fig 2F). Consistent with previous findings (Kassas *et al*, 2017), these data suggest that Opi1 senses not only the PA content, but also a bulk membrane property that is related to the lipid acyl chains in the membrane core. The sensitivity of Opi1 to a collective membrane property is particularly interesting in light of its physiological role. By sensing both PA and the molecular lipid packing density of the ER, Opi1 integrates two crucial signals for the regulation of membrane biogenesis (Hofbauer *et al*, 2014).

### Interfacial hydrophobicity tuning of an AH modulates membrane binding

Having established that Opi1 uses an AH for membrane recognition, we set out to tune its properties by rational design. If the hydrophobic face of the AH contributed to membrane recruitment, an increase of the interfacial hydrophobicity should increase membrane binding, whereas a decrease should lower it. We thus turned our focus on Y127, the residue with the highest interfacial hydrophobicity present in the hydrophobic face of the Opi1 AH (Wimley & White, 1996). A Y127W mutation was generated to increase the hydrophobicity, whereas the mutations Y127A and Y127D were introduced to decrease it (Wimley & White, 1996). As alternative approaches to tune the membrane binding properties of Opi1, we introduced a G120W mutation in the hydrophobic face of the AH to increase both the hydrophobicity and the helical propensity (Wimley & White, 1996; Monné *et al*, 1999). We also increased the length of the AH by a simple duplication of the sequence from residue 114 to residue 131 (2xAH). These variants of MBP-Opi1R^180*^ (Fig 3A) were isolated (Fig 3B) and subjected to membrane binding assays. The proportion of unsaturated lipid acyl chains was kept at 75% in all experiments, while the molar fraction of PA was varied between 0 mol% and 50 mol% (Fig 3C-I). Liposomes containing 40 mol% PS were used to test if changes in the hydrophobic face of the AH and the resulting possible changes in its positioning relative to the bilayer might contribute to the headgroup selectivity of Opi1 (Fig 3J). The binding assays revealed that none of the MBP-Opi1R^180*^ variants binds to liposomes with 5 mol% PA or less (Fig 3C,D). Consistent with earlier findings (Loewen, 2004; Young *et al*, 2010), this suggests that electrostatic interactions between the PA headgroup and basic residues of Opi1 (Fig 1D) are a prerequisite for membrane recruitment. However, the approach of tuning the AH by targeting its hydrophobic face was successful at intermediate PA levels between 10 and 30 mol% (Fig 3E-G): decreasing the interfacial hydrophobicity at position 127 by the Y127D and Y127A mutations weakened membrane binding of Opi1, whereas increasing the hydrophobicity by the Y127W exchange strengthened it (Fig 3E-G). The G120W mutation increases both the hydrophobicity and the helix propensity of the AH, making it even more effective than the Y127W mutation in supporting membrane recruitment of Opi1, as apparent from the experiments with liposomes containing 20 and 30 mol% PA (Fig 3F,G). Not surprisingly, the duplication of the AH turned out to be most effective in supporting membrane recruitment of Opi1 (Fig 3E-G). At higher levels of PA, Opi1 efficiently bound to the liposomes irrespective of the mutations introduced in the hydrophobic face of the AH (Fig 3H,I). This suggests that the electrostatic interactions between the PA headgroup and the hydrophilic face of the AH can become dominant over the contribution of the hydrophobic face in mediating membrane binding. The intriguing selectivity of the Opi1 AH for PA over PS observed in experiments with the synthetic Opi1^111-128^ peptide (Fig 2D) was further corroborated by these experiments designed to tune Opi1 membrane binding propensity. For all of the tested variants of Opi1, the binding to liposomes containing 40 mol% PS (Fig 3J) was weaker than the binding to liposomes containing 20 mol% PA (Fig 3F). From this observation we conclude that the hydrophobic face of the AH is capable to modulate membrane binding at intermediate PA levels but does not critically contribute to the selectivity for PA.

**Figure 3:**
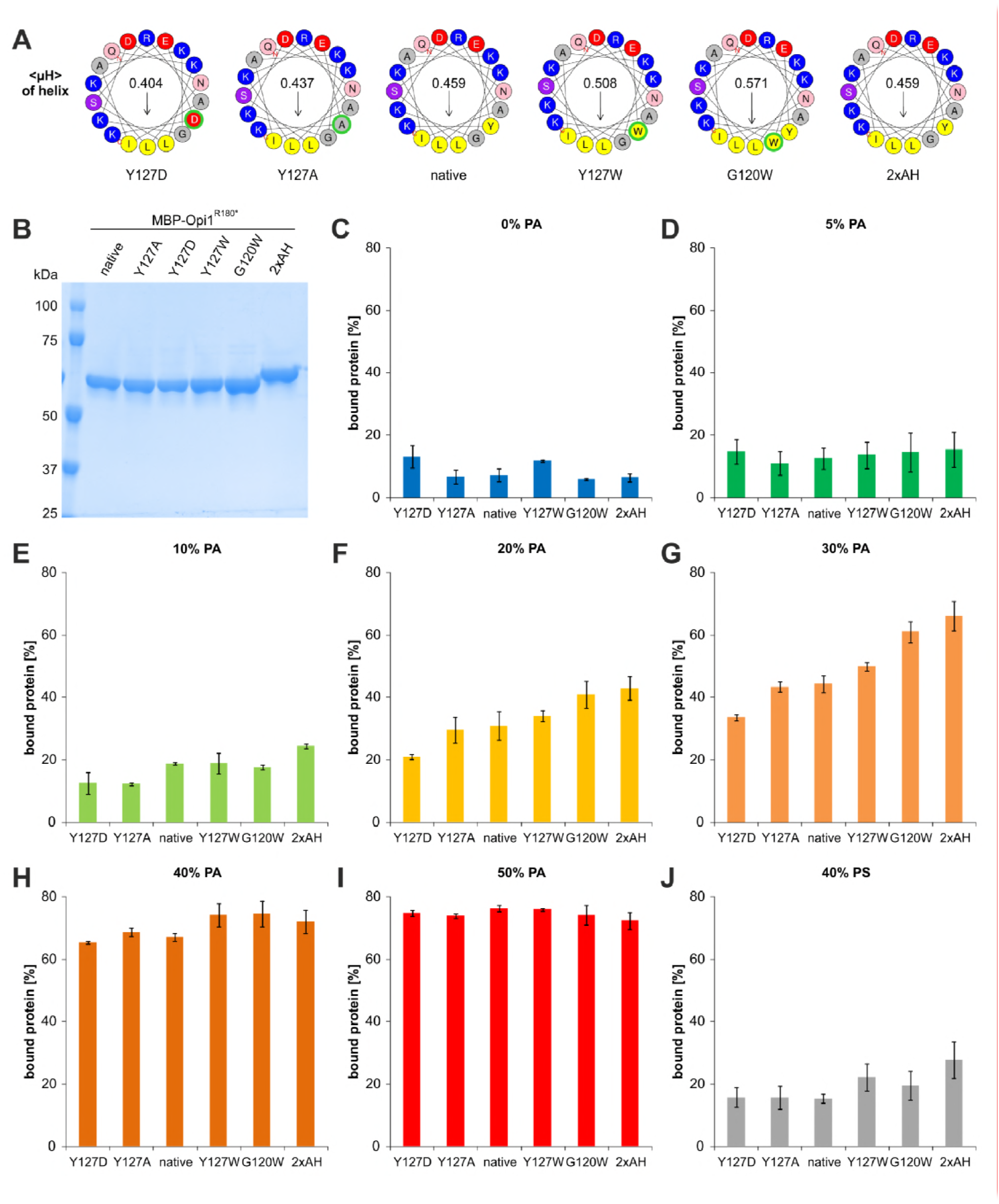
Interfacial hydrophobicity tuning affects membrane binding of MBP-Opi1R^180*^. A Helical wheel representations of different MBP-Opi1R^180*^ variants using the Heliquest tool. B The indicated MBP-Opi1R^180*^ variants were purified by a two-step purification using affinity and size exclusion chromatography (Superdex 200 Increase 10/300 GL), and 1 μg of each protein was subjected to SDS-PAGE for quality control. C-J Opi1 binding assays to liposomes containing increasing proportions of POPA (C-I) or 40 mol% POPS (J) as performed for Figure 2C. The percentage of Opi1 found in the top fraction of the gradient was determined by densiometry. The bar diagrams show the average of 2 independent experiments for C, E, H and I, and 3 independent experiments for D, F, G and J. Values for the native MBP-Opi1R^180*^ variant with 0% PA, 20% PA and 40% PS are the average of 9, 11 and 9 independent experiments, respectively, and are the same as in Figure 2D and Figure 6C. Error bars represent the standard deviation.

### Interfacial hydrophobicity tuning to control cellular signaling

The molar fraction of PA in total cell lipidomes has been reported to lie between 5 and 15 mol% depending on the growth conditions (Klose *et al*, 2012). Other studies reported that PA makes up only 0.2% to 3% of the glycerophospholipids in isolated microsomal fractions from yeast (Zinser *et al*, 1991). Thus, the cellular level of PA is typically much lower than the 10-30 mol% of PA for which interfacial hydrophobicity tuning proved efficient (Fig 3E-G) and the 40-50 mol% of PA (Fig 3H,I) necessary for the quantitative membrane recruitment of Opi1 *in vitro* (Fig 2,3). In cells, however, Opi1 binding is stabilized by the ER membrane-spanning protein Scs2 interacting with the Opi1 FFAT motif (Fig 1C) (Loewen *et al*, 2003; Loewen, 2004). Therefore, it is conceivable that a quantitative recruitment of Opi1 to the ER occurs already at lower levels of PA *in vivo*. The presence of a co-receptor is a complicating factor for the rational design of Opi1 in the physiological context. In order to validate our tuning approach in cells, we generated Opi1-mGFP knock-in constructs that were targeted to the endogenous gene locus of *OPI1*. Notably, the C-terminal tagging of Opi1 does not cause any profound functional defect as previously shown (Gaspar *et al*, 2011). Based on the conclusions from the *in vitro* experiments (Fig 2,3), we generated a series of mutant Opi1-mGFP constructs to manipulate membrane binding properties by tuning the interfacial hydrophobicity of the AH. Aiming at the disruption of the amphipathic character of the PA binding domain, we generated three mutant variants that introduce a charged residue in the interfacial region: Y127R, Y127K, and Y127D. A Y127A mutation was generated to test if the mild membrane binding defect of Opi1 observed *in vitro* (Fig 3E) would be detectable by sensitive cell based assays also *in vivo*. To test if an aromatic residue at position of Y127 is specifically required for normal signaling, we generated a Y127L mutation featuring a similar interfacial hydrophobicity (Wimley & White, 1996), while lacking the aromatic character. The G120W and the G120W/Y127A mutations were generated to cover a broader spectrum of interfacial hydrophobicities in the AH. Finally, the DNLS mutant (Opi1^KRQK->AAQA^) lacking the nuclear localization sequence served as an additional control. These rationally designed mutant variants of Opi1 were subjected to an in depth phenotypic characterization: i) Inositol-auxotrophy on solid media identified mutations that destabilize the binding of Opi1 to the ER (Fig 4A), ii) live cell confocal microscopy provided complementary information on the subcellular localization of Opi1-mGFP variants (Fig 4B), iii) the cellular growth rate in liquid medium during acute inositol depletion provided an indirect, yet quantitative measure for defective membrane recruitment of Opi1 (Fig 4C), and iv) the <u>o</u>ver<u>p</u>roduction of <u>i</u>nositol (Opi^−^) phenotype identified Opi1 mutant variants that are stabilized at the ER, thereby causing a deregulated *INO1* expression and inositol secretion from cells. Secreted inositol can be detected by an inositol auxotrophic tester strain forming a red halo around a spotted, inositol-secreting colony on an otherwise inositol-free plate (Fig 4C *middle panel*) (Greenberg *et al*, 1982). An immunoblot to detect the steady-state level of the generated Opi1-mGFP variants served to complement the phenotypic characterization. As the *OPI1* promoter contains a UAS_INO_ sequence, it is repressed by the Opi1 gene product itself (Schüller *et al*, 1995). Thus, the steady-state level of Opi1-mGFP should correlate with the membrane binding strength of the engineered Opi1 variants (Fig 4C). A decreased level of mutant Opi1-mGFP compared to wild type Opi1-mGFP indicates weaker membrane binding and a repression at the *OPI1* locus, while higher cellular levels indicate enhanced membrane binding of the respective Opi1-mGFP variant. Using this palette of phenotypic readouts, we studied the responsiveness of the Opi1 regulatory system in order to characterize rationally designed AH variants (Fig 4A-C).

**Figure 4:**
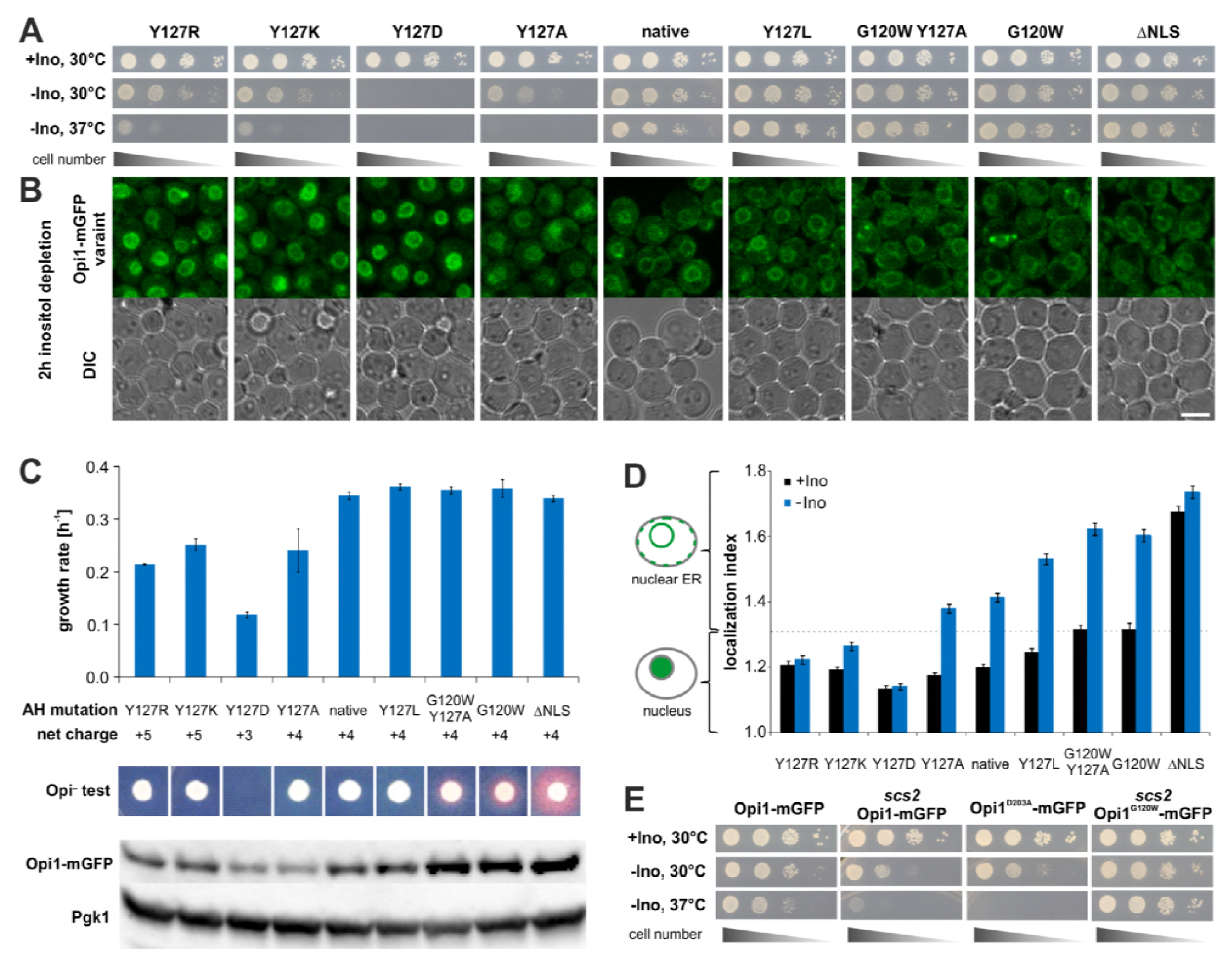
*In vivo* validation of interfacial hydrophobicity tuning. A Viability assays of strains with chromosomally integrated Opi1-mGFP variants. Plates were scanned after 2 days of cultivation on solid media either containing (−Ino) or lacking inositol (−Ino) at the indicated temperatures. The ΔNLS mutant (K109A, R110A, K112A) lacks the nuclear localization sequence. B Representative microscopic images of live cells expressing Opi1-mGFP variants cultivated for two hours after inositol depletion in liquid media. Scale bar, 5 μm. C Phenotypical characterization of the indicated Opi1-mGFP variants. The growth rate in medium lacking inositol between three and six hours of inositol-depletion was determined by absorption spectroscopy. The Opi^−^ phenotype was used to monitor overproduction and secretion of inositol apparent as a red halo from an inositol-auxotrophic tester strain around the spotted colonies. Steady-state levels of the MBP-Opi1R^180*^ variants in the lysates of cells cultivated for two hours in inositol-lacking media were analyzed by immunoblotting. Opi1-mGFP was detected using an anti-GFP antibody with anti-Pgk1 antibody as internal control. D Quantification of microscopic images from B. The localization index is a semi-quantitative measure for the subcellular localization of the indicated Opi1-mGFP variants. A localization index below 1.3 indicates a nuclear localization of the respective Opi1-mGFP variant, while a localization index above 1.3 indicates ER localization. The data points represent the average of four independent replicates with more than 190 analyzed individual cells per strain. The error bars represent the standard deviation. E Viability assay of chromosomally integrated Opi1-mGFP variants in wild type and scs2 deletion strain backgrounds. Plates were scanned after 2 days of growth on solid media at the indicated temperatures.

All mutant variants showed normal growth on solid media plates containing inositol (Fig 4A), whereas the disruption of the amphipathic character by the Y127K, Y127R, and Y127D mutations and a decreased interfacial hydrophobicity by the Y127A mutation caused significant growth defects on media lacking inositol, which were even more pronounced at an elevated growth temperature of 37°C (Fig 4A).

Consistently, the localization of mutant Opi1-mGFP was greatly affected during acute inositol depletion, which is known to cause an increase of the cellular level of PA (Fig 4B) (Hofbauer *et al*, 2014). While the native Opi1-mGFP localized to the ER under these conditions, the Y127K, Y127R, Y127D and less so the Y127A showed an increased nuclear localization, suggesting a failure of these mutants to bind PA at the ER membrane. As these Opi1-mGFP variants cannot activate membrane biogenesis even during a build-up of PA, the respective cells show profound growth defects during prolonged inositol depletion between three and six hours (Fig 4C). Immunoblotting for Opi1-mGFP from cellular lysates revealed that the steady-state level of the Y127R, Y127K, Y127D and the Y127A variants were reduced compared to native Opi1-mGFP (Fig 4C). This suggests that the increased nuclear localization of these mutants represses the *OPI1* gene, resulting in decreased steady-state protein levels. Thus, all mutants disrupting the AH and lowering its interfacial hydrophobicity also disrupt Opi1 binding to the ER and impose profound growth defects during inositol depletion. Intriguingly, the substitution of Y127 with basic residues (Y127R and Y127K) caused milder growth defects than the Y127D mutation (Fig 4A,C). This may indicate that basic side chains present at the hydrophobic face of the AH can ‘snorkel’ towards the aqueous environment (Öjemalm *et al*, 2016) and rescue defective membrane recruitment of the protein by providing additional electrostatic interactions with PA lipid headgroups.

The rational design of the AH also led to Opi1-mGFP variants with enhanced membrane binding properties. Whereas the Y127L variant appeared most similar to native Opi1-mGFP in all assays, the G120W/Y127A and the G120W variants were better binders. Cells expressing these mutant variants did neither exhibit growth defects in inositol-lacking media (Fig 4A,C), nor did localization of Opi1 respond in any way during acute inositol depletion (Fig 4B). Instead, the increased ER membrane binding of these mutants led to Opi^−^ phenotypes and the secretion of inositol (Fig 4C). The most pronounced Opi^−^ phenotype was observed for the ΔNLS variant that cannot enter the nucleus due to the absence of the nuclear localization sequence (Fig 4C). Consistent with previous findings, this suggests a dynamic equilibrium of Opi1 between two pools, at the ER and Opi1 in the nucleus (Loewen, 2004). The increased binding to the ER membrane by the G120W/Y127A, G120W, and ΔNLS variants also caused a de-repression of *OPI1* gene expression in the nucleus, which ultimately resulted in increased steady-state levels of these mutant variants (Fig 4C) that reside in the ER (Fig 4B). Together, these data provide compelling evidence that a rational design of interfacial hydrophobicity is suitable to tune membrane binding *in vitro* and subcellular localization of Opi1 and its regulatory function *in vivo*.

Using an automated script to analyze microscopic images taken from the entire set of Opi1-mGFP variants before and during acute inositol depletion (as in Fig 4B), we further quantitatively determined the impact of the AH on the subcellular localization. We established a localization index based on the shape and size of the GFP-positive signal in cells that allowed us to quantitatively assess the impact of interfacial hydrophobicity tuning on the subcellular localization of Opi1-mGFP. The automated analysis validated our conclusions from the visual inspection (Fig 4B) and established a ranking of the disruptive mutations. The binding of Opi1 variants to the ER increases in the order Y127D<Y127R<Y127K<Y127A<native and the stabilizing variants can be ranked from weaker binders to better binders as native<Y127L<G120W/Y120A~G120W (Fig 4D). The analysis further revealed that an aromatic residue at position 127 is not required for effective membrane binding as Y127 can be substituted with leucine without disrupting membrane binding. Tuning the interfacial hydrophobicity by other mutations clearly affected the subcellular localization of Opi1.

Since Opi1 is a powerful transcriptional repressor that regulates lipid metabolism and membrane biogenesis we set out to analyze whether the aberrant localizations of the engineered Opi1 variants are due to secondary effects, e.g. an altered ER lipid composition. To test for this, we constructed heterozygous diploid yeast cells bearing one allele encoding for a mutant variant of Opi1-mGFP and one allele encoding for the native Opi1 fused to red fluorescent protein (Opi1-mRFP) (Fig EV3). Consistent with the analyses of haploid cells, the subcellular localization of Opi1-mGFP was greatly affected by mutations in the AH (Fig EV3), whereas native Opi1-mRFP remained predominantly associated with the ER membrane irrespective of the co-expressed Opi1-mGFP mutant variant (Fig EV3). Even though the mutant and the native variant of Opi1 were facing the same ER membrane, they exhibited striking differences in their subcellular distribution. This suggests that the mutations affect the membrane recognition *per se* and argues against a perturbed membrane lipid composition or other secondary effects as a cause for Opi1 mislocalization.

Together, these data establish the tuning of interfacial hydrophobicity as a promising tool for custom-designing membrane recruitment of proteins. In order to challenge this proposal, we tested if a rationally designed Opi1^G120W^-mGFP variant with an increased membrane binding potential (Fig 3F,G) could compensate for a loss of Scs2, the co-receptor for Opi1 at the ER-membrane. The loss of Scs2 causes significant growth defects on media lacking inositol, especially at elevated temperatures (Loewen et al, 2003; Gaspar et al, 2017) (Fig 4E). These effects are phenocopied by a mutation in the Opi1 FFAT motif (Opi1D203A-mGFP) that disrupts the interaction between Opi1 with Scs2 (Fig 5E; Loewen and Levine, 2005). Strikingly, the G120W mutation in Opi1 is sufficient to rescue the inositol auxotrophy associated with a loss of Scs2, thus providing further evidence that tuning of the interfacial hydrophobicity provides a powerful tool to manipulate the membrane binding of AH containing proteins: the exchange of a single residue in the AH compensates for the loss of the co-receptor Scs2 in the ER membrane.

**Figure 5:**
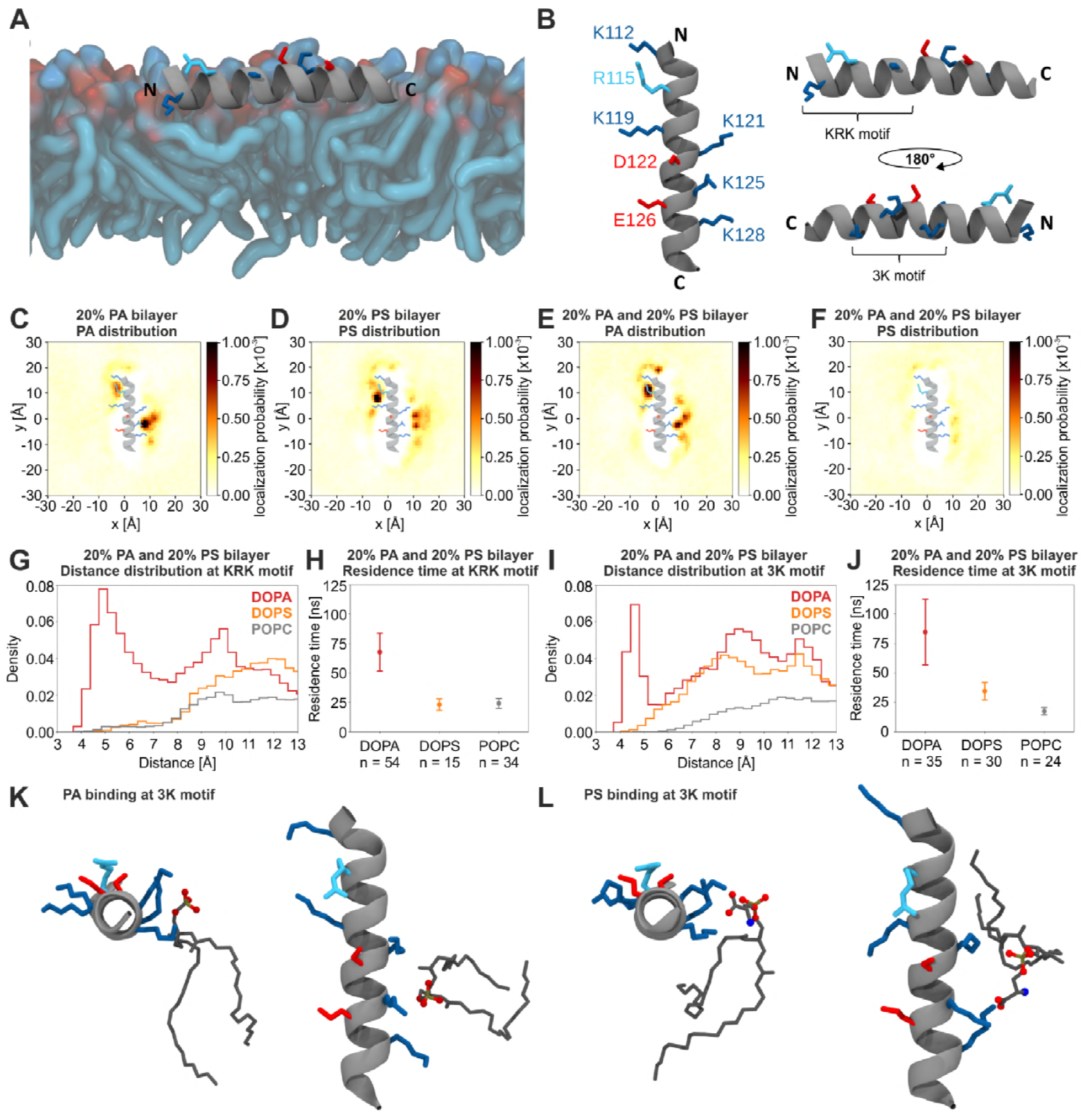
Molecular dynamics simulations of PA and PS binding to Opi1 AH. A Representative structure of the AH of Opi1^120-132^ in a lipid bilayer from an all-atom MD simulation. The helix is represented as a gray ribbon with charged amino acids lysine (blue), arginine (cyan), aspartate and glutamate (red) as sticks. The AH remained folded and inserted in the membrane over the course of >5us. Lipids are shown as tubes (carbon, blue; oxygen, red). Only the top membrane leaflet is shown. B Representative structure of Opi1 AH (Opi1^120-132^) shown from three different directions: top (left panel), N- to C-terminus (top right), and C- to N-terminus (bottom right). The two lipid binding sites KRK- and 3K-motif are annotated. Color code as in A. C-F Time-averaged positions of phosphate head groups from two lipid species (C and E: DOPA, D and F: DOPS) in three different all-atom MD simulations (C: 20% PA, D: 20% PS, E and F: mixed 20% PA / 20% PS). Colors indicate the probability to observe a given lipid at that position over the course of the trajectory. DOPA lipids localize closer to the 3K-motif (C) than DOPS lipids (D). DOPA displaces DOPS in a mixed bilayer at both motifs (E and F). The Opi1 AH was superimposed to show the localization of high density regions around positively charged residues (see 5B for annotations). G,I Distribution of pairwise distances calculated from a binding motif and all lipids of a given species in a mixed bilayer. DOPA (red) fully displaces DOPS (orange) and POPC (gray) from the KRK-motif (5G). At the 3K-motif DOPA is closest to the AH, while DOPS is found at a higher distance (5I). The binding motif was defined as the combined backbone Cα center of mass of three amino acids (5G: K112, R115, and K119 or 5I: K121, K125, and K128). H,J Residence times of different lipid species at the two motifs KRK (5H) and 3K (5J) for the three lipids DOPA (red), DOPS (orange), and POPC (gray). DOPA shows a 3-fold higher residence time than DOPS and POPC. The number of observed binding events is indicated as a label on the x-axis. K,L Representative structures of a DOPA (K) or DOPS (L) lipid interacting with lysines at the 3K-motif shown from the C-terminus (left panels, K and L) and top (right panels, K and L). DOPA interacts with oxygens at the phosphate head group. DOPS additionally interacts with the carbonyl oxygen of serine, keeping it at a higher distance from the AH. Amino acids are colored the same way as in 5A.

### Identification of a structural motif for PA-selectivity

Both Opi1 and its isolated AH (Opi1^111-128^) exhibit a striking selectivity for PA over PS *in vitro* (Fig 2,3). We used multi-resolution molecular dynamics (MD) simulations to gain insight into the mechanism of selective PA recognition. We modeled an Opi1^111-132^ peptide as an alpha-helix and created two different systems including the AH, solvent, and lipid bilayers, containing either 20 mol% PA or 20 mol% PS, for coarse-grained (CG) MD simulations using the MARTINI representation (Marrink *et al*, 2007; De Jong *et al*, 2013). Both lipid bilayers were composed of 60 mol% POPC, 20 mol% DOPC and either 20 mol% DOPA or 20 mol% DOPS to yield bilayers with 70% monounsaturated acyl chains. After initial CG simulations allowing the Opi1 peptide to associate spontaneously and equilibrate with the membrane, each system was lifted from the CG representation to a fully atomistic representation in the CHARMM36 force field (Jo *et al*, 2007, 2008, 2009, 2014; Lee *et al*, 2016; Wassenaar *et al*, 2014). We simulated both the PA and PS membrane systems for 5 μs, during which the AH remained stably associated with the respective lipid bilayer (Fig 5A,B). All lipid species were mobile and exchanged between regions close to the AH peptide and the bulk membrane. The hydrophobic residues of the AH pointed into the hydrophobic core of the bilayer, while the hydrophilic residues were situated in the membrane-water interfacial (Fig 5 A,B).

We identified structural features of particular importance for the binding of PA and PS from the localization probability of these lipids relative to the AH peptide (Fig 5C,D). We found that the AH is capable to attract and enrich both PA and PS in its vicinity (Fig 5C,D), but neither DOPC nor POPC (Fig EV4A,B). This local enrichment of anionic lipids occurred only in the membrane leaflet containing the AH peptide and was absent in the opposing leaflet of the lipid bilayer (Fig EV4A,B). Most strikingly, PA and PS were concentrated at discrete hot-spots of binding, in the proximity of the ‘KRK motif’ formed by K112, R115 and K119, and the ‘3K motif’ formed by K121, K125 and K128 pointing in the opposite direction from the AH (Fig 5B). The PA and PS binding hotspots differed in their position and distance relative to the KRK and 3K motifs. These differences are not due to different charges in the lipid headgroup region, since the net charge of PA and PS lipids was maintained at -1 throughout the entire atomistic MD simulation. Thus, MD simulations provide a first-time evidence for a different mode of PA and PS binding by the AH of Opi1.

In order to further characterize these different modes of binding, we prepared an MD simulation with a mixed bilayer containing both 20 mol% PA and 20 mol% PS with the headgroup charges maintained at -1. Strikingly, the binding of PA to the KRK and the 3K motif prevented an accumulation of PS in the vicinity of the AH (Fig 5 E,F), suggesting that PA binding dominates over PS binding. We next analyzed the distance distribution of PA and PS relative to the KRK (Fig 5G) and the 3K motif (Fig 5I). The distance profile revealed a population of PA lipids interacting intimately with the 3K motif at distances between 4 and 6 Å (Fig 5G,I). Neither PS (Fig 5G,I) nor PC (Fig EV4D-E) lipids formed such a population and were found only at greater distances, thereby suggesting a specific mode of interaction between the two binding motifs and PA. This interpretation was corroborated by a quantitative analysis of the lipid residence times in the proximity of both motifs (Fig 5H,J). PA lipids dwell significantly longer close to the KRK and the 3K motifs than PS or PC lipids (Fig 5H,J). Our atomistic MD simulations therefore suggest that the AH of Opi1 interacts more intimately and stably with PA than with PS or PC lipids. A closer inspection of the trajectories revealed that the phosphate moiety of PA can be enwrapped by the three basic residues of the 3K motif forming a three-finger grip (Fig 5K), which fail to accommodate the larger PS headgroup (Fig 5L). It is tempting to speculate that the striking selectivity of Opi1 for PA lipids observed *in vitro* (Fig 2,3) is mediated by PA-selective three-finger grips.

### Tuning lipid selectivity by rational design

Having identified a putative mechanism for PA selectivity by MD simulations, we set out to validate this finding experimentally. The AH of Opi1 contains two acidic residues and several basic residues possibly forming two PA-selective three-finger grips (Fig 5B). We wondered, whether PA selectivity may be assisted by the specific arrangement of attractive and repelling forces brought about by basic and acidic residues in the hydrophilic face of the AH. As an alternative but not mutually exclusive model, we speculated that a PA-selective three-finger grip, which relies on just three basic residues on one side of an AH, may have specific structural requirements for efficiently enwrapping the PA headgroup. Thus, the substitution of the lysine-rich AH to an all-arginine AH was expected to affect PA selectivity resulting from the steric differences between arginine and lysine side chains that might prevent the formation of a tight-fitting three-finger grip. Moreover, because the guanidinium group of arginine can form multiple hydrogen bonds with water and the polar lipid headgroups (Li *et al*, 2013), we reasoned that an all-arginine AH might not be able to efficiently distinguish between PA and other anionic glycerophospholipids. We thus generated an MBP-Opi1R^180*^ D122K/E126K variant, to increase the net charge of the AH from +4 to +8, and a second variant in which all lysine residues of the AH were replaced with arginines (5Kà5R), therefore not changing the net charge of the AH (Fig 6A).

**Figure 6:**
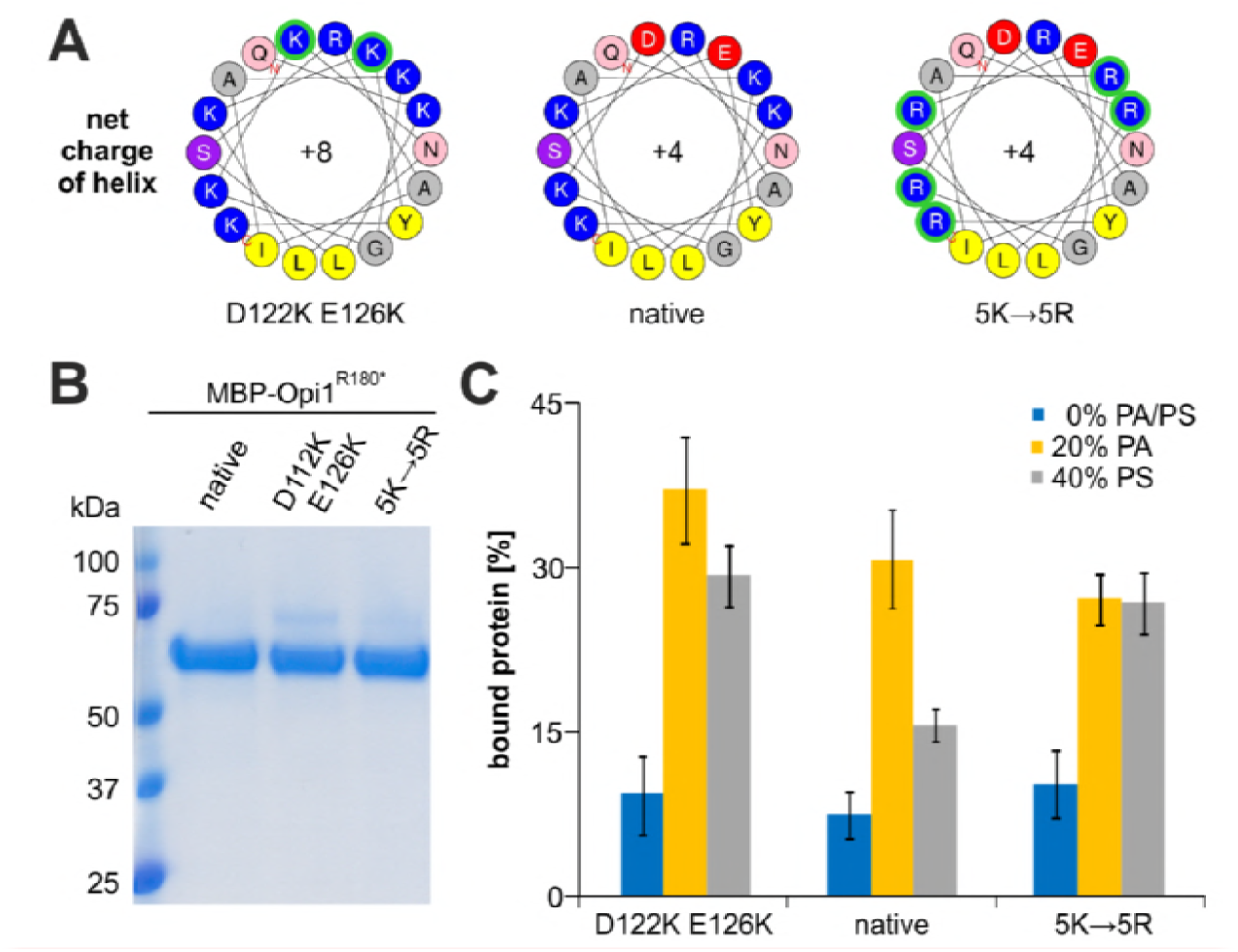
The hydrophilic face of Opi1´s AH is crucial for lipid headgroup selectivity. A Helical wheel representation of MBP-Opi1R^180*^ variants using the Heliquest tool. B The indicated MBP-Opi1R^180*^ variants were purified by a two-step purification using affinity and size exclusion chromatography (Superdex 200 Increase 10/300 GL), and 1 μg of each protein was subjected to SDS-PAGE for quality control. C Liposome binding assays with the indicated MBP-Opi1R^180*^ variants were performed as in Figure 2C. The percentage of Opi1 found in the top fraction of the gradient was determined by densiometry. The bar diagrams show the average of 5 independent experiments each for the D112K E126K mutant and 3 independent experiments each for the 5Kà5R mutant. Values for the native MBP-Opi1R^180*^ variant with 0% PA, 20% PA and 40% PS are the average of 9, 11 and 9 independent experiments, respectively, and are the same as in Figure 2D and in Figure 3C,F,J. Error bars represent the standard deviation.

After isolation of the indicated MBP-Opi1R^180*^ variants by affinity purification and size exclusion chromatography (Fig 6B), we characterized their binding to PA- and PS-containing liposomes using liposome flotation assays (Fig 6C). The PA-containing liposomes were composed of 50 mol% DOPC, 30 mol% POPC and 20 mol% POPA, while PS-containing liposomes were composed of 50 mol% DOPC, 10 mol% POPC and 40 mol% POPS to compensate for the possible charge differences in the headgroup region. Liposomes composed solely of 50 mol% DOPC and 50 mol% POPC served as negative control (Fig 6C). The D122K E126K mutant with a net charge of +8 showed an increased membrane binding to both PA- and PS-containing liposomes, suggesting that increased electrostatic interactions between the AH and anionic lipids supports membrane binding and that the acidic residues have only a minor – if any – function for PA selectivity. The 5Kà5R mutant completely lost the preference for PA- over PS-containing liposomes, confirming our hypothesis that the three-finger grips based on lysine-rich motifs in the AH of Opi1 represent crucial structural elements for the intriguing PA-selectivity of Opi1.

## Discussion

Cells must adjust their lipid metabolism to the available pool of nutrients in order to maintain membrane integrity in response to metabolic challenge or environmental cues. Membrane biogenesis relies on the coordinated regulation of lipid biosynthetic genes and enzymes and is under tight control of the transcriptional repressor Opi1 in *S. cerevisiae* (Henry *et al*, 2012; Schuck *et al*, 2009). The identification and validation of an AH in Opi1 (Fig 1-3) provides a conceptual framework for the previous observations that Opi1 binding to the ER is controlled by the abundance of PA (Loewen, 2004), the lipid headgroup composition (Young *et al*, 2010; Putta *et al*, 2016), the intracellular pH (Young *et al*, 2010), the acyl chain length (Hofbauer *et al*, 2014), the degree of lipid unsaturation (Fig 2F) (Kassas *et al*, 2017), and membrane curvature (Fig 2F) (Kassas *et al*, 2017). Analogous to ALPS motif containing proteins such as GMAP-210 or ArfGAP (Antonny, 2011), Opi1 senses the molecular packing density of lipids by folding an AH into the ER membrane. This PA-selective AH facilitates the Opi1 regulatory circuit to integrate two crucial metabolic input parameters: the PA-abundance and the lipid packing density as a proxy for the lipid acyl chain composition. This feature may have a crucial role in maintaining membrane homeostasis under conditions of ER stress. It is known that aberrant lipid compositions of the ER membrane can cause ER stress and activate of the unfolded protein response (UPR) (Halbleib *et al*, 2017; Walter & Ron, 2011). The UPR controls a large transcriptional program regulating the protein folding capacity of the ER, the ER-associated degradation (ERAD) machinery, the capacity of the secretory pathway, as well as membrane biogenesis and ER abundance (Travers *et al*, 2000; Walter & Ron, 2011). Originally identified as a stress response to accumulating unfolded proteins in the lumen of the ER, it is now clear that the UPR is directly activated also by aberrant lipid compositions (i.e. lipid bilayer stress) (Halbleib *et al*, 2017). Perhaps the most prominent condition of lipid bilayer stress is an increased proportion of saturated lipids (Surma *et al*, 2013; Kitai *et al*, 2013). Under this condition, however, the activation of the UPR initiates a detrimental, positive feedback loop. Enforced membrane biogenesis controlled by the UPR causes a more rapid consumption of coenzyme A (CoA)-activated fatty acids in glycerophospholipids, thereby competing with the desaturase Ole1 to oxidize saturated to unsaturated fatty acids (Ballweg & Ernst, 2017). Thus, the activation of the UPR by saturated membrane lipids causes the biosynthesis of even more saturated lipids, inducing more ER-stress and leading to severe morphological changes of the ER and other organelles (Surma *et al*, 2013; Schneiter & Kohlwein, 1997). By sensing the molecular lipid packing density of the ER membrane, Opi1 provides a means to interrupt this vicious circle. We have shown that increased lipid packing densities destabilize the binding of MBP-Opi1^180*^ to PA-containing liposomes (Fig 2F). In the analogous *in vivo* situation, this would put a break on membrane biogenesis and interrupt the vicious circle induced by saturated lipids. It is tempting to speculate that Opi1 and the inositol-requiring enzyme Ire1, the most ancient and conserved UPR transducer, are part of an integrated regulatory circuit to control ER abundance. The fact that inositol is a key regulator of both Ire1 and Opi1 might be more than coincidence, pointing to a possible auto-regulatory crosstalk between two major regulatory pathways of membrane biogenesis. This view is consistent with previous observations that Opi1 and Ire1 jointly regulate the abundance of the ER and that loss of Opi1 alleviates ER-stress by increasing membrane biogenesis (Velázquez *et al*, 2016; Schuck *et al*, 2009).

Similarly to ALPS motif containing proteins (Drin & Antonny, 2010), Opi1 uses an AH that folds into the cytosolic membrane leaflet of the ER. Our experiments show that the strength of membrane binding is largely determined by the interplay of electrostatic and hydrophobic interactions. In contrast to classical ALPS motifs featuring small polar residues on the hydrophilic face of the AH (Drin & Antonny, 2010), the AH of Opi1 is characterized by several positively charged residues on the hydrophilic face. Electrostatic interactions between these residues and the anionic PA headgroup are dominant determinants for membrane binding and lipid selectivity (Fig 3). Nevertheless, the hydrophobic portion of the AH substantially modulates the strength of membrane binding at intermediate concentrations of PA *in vitro* as well as *in vivo* (Fig 3,4).

Using the Opi1 regulatory system as a phenotypic show-box, we established interfacial hydrophobicity tuning as an approach to rationally design the membrane binding strength of Opi1. This intuitive and straight-forward strategy is likely applicable to other membrane binding AHs that are frequently found in enzymes and regulators of lipid metabolism (Puth *et al*, 2015), tethering proteins of the secretory pathway (Magdeleine *et al*, 2016), and in antimicrobial peptides (Shai, 1999). The modulatory potential of interfacial hydrophobicity tuning is best illustrated by the point mutation G120W in Opi1, which is sufficient to compensate for the loss of the Opi1 co-receptor Scs2 in the ER membrane (Fig 4E). We conclude that interfacial hydrophobicity tuning provides a powerful means to design the membrane binding properties of Opi1 or other proteins.

The remarkable selectivity of the Opi1 AH for PA lipids directly raises the question on how this selectivity is achieved; in contrast to the phosphoesters in other glycerophospholipids such as PC, PE, PI and PS, which can be deprotonated only once, the phosphomonoester of PA can be deprotonated twice, yielding ionization states of either -1 or -2 (Kooijman *et al*, 2005). The pKa for the second deprotonation of PA is close to the physiological pH and greatly affected by the membrane context – a phenomenon described by the ‘electrostatic/hydrogen-bond switch’ model (Kooijman *et al*, 2005; Young *et al*, 2010; Shin & Loewen, 2011). When correcting for the different charges in the headgroup region, the plasma membrane sensor protein Spo20, for example, binds PA, PS and PIPs with comparable efficiencies (Horchani *et al*, 2014). In contrast, when compensating for the maximal possible charge difference of PA and PS, Opi1 still exhibited an inherent selectivity for PA-containing membranes (Fig 2D; Fig 3F,J). While the binding of Opi1 to PA is dominated by electrostatics, the headgroup selectivity must be accomplished by other means.

Large numbers of PA- and PS-binding proteins have been identified and yet, no clear signature motif or structural element for the binding of PA lipids has emerged (Stace & Ktistakis, 2006). Here, we identify two sequence elements (KhXRhhhK and KXXhKXXK, where X stands for any and h for an hydrophobic amino acid) in the AH of Opi1 (Fig 1,2) that selectively recognize PA when situated in an AH by forming three-finger grips (Fig 5). This simple structural element based on a linear sequence motif in a defined helical arrangement is likely to exist in many other proteins, thereby paving the way towards identifying and characterizating potential PA-selective functions of these proteins. The protein-tyrosine phosphatase SHP-1, for example, is activated by PA but not PS (Frank *et al*, 1999) and does indeed contain three-finger grip motifs. Our MD simulations have established molecular and dynamic details of PA recognition by three-finger grips: PA is tightly enwrapped by three basic residues and binds more stably than PC or PS (Fig 5H,J). The loss of selectivity upon the substitution of lysine to arginine residues suggests specific steric constraints that may be required for establishing a PA binding three-finger grip (Fig 6). Consistent with this, the membrane-binding AH of Spo20, which features a histidine-rich polar face, binds to anionic lipids, but does not exhibit a particular selectivity for PA *in vitro* (Horchani *et al*, 2014). Our findings therefore provide a theoretical framework for the observations that Opi1, but not Spo20, localizes to PA-enriched subdomains of the nuclear ER upon lipid droplet formation (Ganesan *et al*, 2015; Wolinski *et al*, 2015).

The sensitivity of Opi1 to a collective membrane property is particularly interesting in light of its physiological role. By sensing both PA and the molecular lipid packing density of the ER, Opi1 integrates two crucial signals for the regulation of membrane biogenesis (Hofbauer et al, 2014). When saturated membrane lipids accumulate to an extent that causes ER-stress (Surma et al, 2013; Halbleib et al, 2017; Pineau et al, 2009), the binding of Opi1 to PA at the ER-membrane would be destabilized. Once released from the membrane, Opi1 would translocate to the nucleus and dampen membrane lipid biosynthesis to prevent the production of even more saturated lipids. In this way, Opi1 counteracts lipid bilayer stress caused by saturated membrane lipids. The identification of headgroup selectivity in AHs thus has important implications for the regulation of lipid metabolism in yeast and adds significantly to our understanding of highly specific subcellular membrane targeting and metabolic regulation by AH-containing proteins.

## Acknowledgements

The pMAL-OPI1 plasmid was kindly provided by Dr. George Carman (Rutgers University, New Brunswick, NJ, USA). The pFA6a-mGFP-NatMX6 and pFA6a-mRFP-KanMX4 plasmids were kindly provided by Dr. Klaus Natter (University of Graz, Austria). The AID strain was kindly provided by Dr. Susan A Henry (Cornell University, Ithaca, NY, USA). We thank Dr. Sepp D. Kohlwein for critically reading the manuscript. H.F.H: is funded by the Erwin Schrödinger fellowship of the Austrian Science Fund (FWF): J 3987-B21. The work was supported by the Deutsche Forschungsgemeinschaft (EN608/2-1 to R.E.; SFB807 Transport and Communication across Biological Membranes to R.E. and G.H.; CEF-MC II to E.S. and R.E.; EC-115 to E.S.). M.G., R.C., and G.H. acknowledge support by the Max Planck Society, and discussions with Drs. Lukas Stelzl, Ahmadreza Mehdipour, and Max Linke.

## Author contributions

R.E. supervised the project. H.F.H., M.G., R.C. and R.E. conceived the experimental design. H.F.H. and R.E. wrote the manuscript. H.F.H performed the experimental work. M.G., R.C., and G.H. designed and conducted MD simulations, and analyzed the data. S.C.F. and E.S. supported quantitative live cell microscopy. A.S., and A.F. performed cryo-electron microscopic imaging of liposomes.

## Materials and Methods

### Reagents

If not stated otherwise, all reagents used in this study were of analytic grade and purchased from Sigma Aldrich or Roth. Following lipids were purchased from Avanti Polar Lipids: 1,2-dioleoyl-*sn*-glycero-3-phosphate (DOPA; 840875), 1-palmitoyl-2-oleoyl-*sn*-glycero-3-phosphate (POPA; 840857), 1,2-dioleoyl-*sn*-glycero-3-phosphocholine (DOPC; 850375), 1-palmitoyl-2-oleoyl-*sn*-glycero-3-phosphocholine (POPC; 850457) and 1-palmitoyl-2-oleoyl-*sn*-glycero-3-phospho-L-serine (POPS; 840034). The amylose resin was purchased from New England Biolabs, while the yeast nitrogen base (w/o amino acids and inositol), the complete supplement mixture, and agar were purchased from ForMedium.

*Escherichia coli and Saccharomyces cerevisiae* strains used in this study are listed in Table EV1, plasmids are listed in Table EV2 and primers are listed in Table EV3.

### Peptide synthesis

The synthetic, biotinylated Opi1^111-128^ peptide (Biotin-QKLSRAIAKGKDNLKEYK-CONH2; MW 2315.87 g/mol) was purchased from ZIK B CUBE (TU Dresden) and purified to >90% purity. The quality of the isolated peptide was validated by the supplier using mass spectrometry. The peptide was dissolved in NaPi buffer (20 mM sodium phosphate, 150 ml NaCl, pH 7.4) to yield a stock concentration of 10 mg/ml (4.3 mM) for further analyses.

### Preparation and characterization of synthetic liposomes

Glycerophospholipid powders were dissolved in chloroform to yield 20 mg/ml stocks. The desired lipid compositions were mixed in 15 ml Pyrex glass tubes, dried under a steam of nitrogen, and put in an exsiccator over night to quantitatively remove the residual organic solvent. Lipid films were rehydrated for 2 hours at room temperature using NaPi buffer (20 mM sodium phosphate, 150 mM NaCl, pH 7.4) to a final lipid concentration of 4 mM and mixed every 15 min. After rehydration, liposomes were subjected to 5 freeze/thaw cycles using liquid nitrogen and a 50°C heating block.

Stepwise extrusion of these liposomes using the LipoFast^TM^ extruder (Avestin) with 21 passages through pore size filters of 200 nm, 100 nm, 50 nm and 30 nm (Avanti Polar Lipids) yielded monodisperse populations of mostly unilamellar vesicles. The average size of the liposomes was determined using a NanoSight LM10 Nanoparticle Analysis System (Malvern Instruments). The measurements all contained at least 2000 valid tracks and the polydispersity index (PdI) was below 0.1 in all cases. The quality of the liposome preparation was further analyzed using cryo-electron microscopy. For plunge-freezing, 3.5 μl of the sample were pipetted onto a glow-discharged Quantifoil 3.5/1 grid (Quantifoil) and vitrified using a Vitrobot (FEI) operated at a -1 offset and 4 sec blotting time with pre-wetted filter paper (grade 595; Whatman). Cryo-grids were imaged with a Tecnai F-30 (FEI) electron microscope operated at 300 kV. Images were recorded on a US4000 CCD camera (Gatan).

### Circular dichroism spectroscopy

Changes of the secondary structure of the synthetic Opi1^111-128^ peptide was measured using a Jasco J-810 spectropolarimeter (Jasco). 20 mM SDS, 20 mM DDM or 4 mM liposomes in NaPi buffer were mixed with 20 μM peptide in a volume of 200 μl and analyzed in a cuvette with 1 mm path length from 260-190 nm at 22°C. The parameters were as follows: standard sensitivity, 1 nm data pitch, digital integration time of 1 sec, 1 nm bandwidth, 100 nm/min scanning speed and 3 repeats for each measurement. Blank measurements without peptide were subtracted from the spectra prior to analysis. Values below 200 nm were omitted from the final graph due to exceeded high tension voltage (>700 V) of the detector. Mean residue ellipticity (MRE) values were calculated using the formula

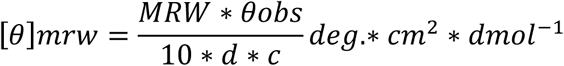

with *MRW* as mean residue weight, *d* as path length in cm and *c* as peptide concentration in g/ml.

### Generation of MBP-Opi1 constructs for expression in *E. coli*

For the expression of the N-terminally tagged MBP-Opi1 fusion protein, the MBP-Opi1R^180*^, MBP-Opi1^S114*^, and mutant variants of MBP-Opi1R^180*^, we used the plasmid pMAL-OPI1 (Sreenivas & Carman, 2003) kindly provided by Dr. George Carman (Rutgers University, NJ, USA). The codons that were mutated and the corresponding primers are listed in Table EV3. Site-directed mutagenesis was either conducted using a slightly modified QuikChange^®^ method (Stratagene) with the PHUSION polymerase (NEB) or using the Q5 site-directed mutagenesis kit (NEB), indicated by a Q5-suffix in the primer names in Table EV3.

### Generation of Opi1-mGFP constructs and chromosomal integration in yeast

For the generation of Opi1-mGFP knock-in constructs, we used the pFA6a-mGFP-NatMX6 construct kindly provided by Dr. Klaus Natter (University of Graz, Austria). First, we amplified a C-terminal *OPI1* homology region that is 3-556 bp downstream from the *OPI1* stop codon with overhangs for *EcoRI* and *SpeI* using the primers RE529 and RE530 to generate plasmid pFA6a-mGFP-NatMX6-C-term-homology by restriction-based cloning. For the generation of pFA6a-promoter-OPI1-mGFP-NatMX6-C-term-homology, we amplified the *OPI1* open reading frame including 275 bases upstream the start codon from genomic DNA with overhangs for the restriction enzymes *HindIII* and *BamHI* using the primers RE528 and RE440 and classical cloning. Site-directed mutagenesis was performed with primers listed in Table EV3. After plasmid isolation and validation by DNA sequencing (SeqLab), 3 μg of the plasmid were digested with *NotI* (NEB) to yield a linear DNA fragment containing the *OPI1* promoter, the open reading frame of *OPI1* and a C-terminal homology region for transforming BY4742 or *scs2* yeast strains using the LiAc method. Transformants were selected on YPD plates containing 100 μg/ml nourseothricin (NTC) and positive clones were verified via DNA sequencing after colony PCR using the primers RE548 and RE916.

For the generation of Opi1-mRFP knock-in constructs, we used the pFA6a-mRFP-KanMX4 construct kindly provided by Dr. Klaus Natter (University of Graz, Austria). The mRFP-KanMX4 cassette was amplified with overhangs for chromosomal insertion upstream of the stop codon of *OPI1* using the primers RE647 and RE648. The amplified fragment was purified using the PCR purification kit (Qiagen) and 1 μg of the purified fragment was used for transformation of BY4741 using the LiAc method. Transformants were selected on YPD plates containing 100 μg/ml kanamycin (KAN) and the correct insertion of mRFP was validated by colony PCR using the primers RE649 and RE646.

Diploid strains bearing Opi1-mGFP as one allele and native Opi1-mRFP as the other allele were generated by mating of the haploid strains followed by selection for diploid cells on YPD medium containing 100 μg/ml NTC and 100 μg/ml KAN.

### Expression and purification of recombinant MBP-Opi1 protein variants

The *E. coli* BL21-CodonPlus(DE3)-RIL strain (Agilent Technologies) was used for heterologous production of MBP-Opi1 variants. Cells were cultivated over night at 37°C under constant agitation (at 220 rpm) in 5 ml lysogeny broth (LB)-medium (1% peptone, 0.5% yeast extract, 1% sodium chloride) containing 100 μg/ml ampicillin (Amp) and 34 μg/ml chloramphenicol (CA). The main culture was inoculated from this over night culture to an optical density (OD_600_) of 0.25 in 250 ml fresh LB Amp/CA medium. At an OD_600_ of 0.5-0.6, heterologous gene expression was induced using 500 μM isopropyl-β-D-thiogalactopyranoside (IPTG). Cells were harvested three hours after induction. The cell pellets (400-500 OD_600_ units) were washed once in ice-cold protein purification (PP) buffer (50 mM HEPES, 150 mM NaCl, 1 mM EDTA, pH 7.4) and stored at -20°C.

Cells were thawed on ice and resuspended in 22 ml ice-cold protein lysis buffer (50 mM HEPES, 150 mM NaCl, 1 mM EDTA, 2 mM DTT, 1 mM AEBSF, 10 μg/ml chymostatin, 10 μg/ml antipain, 10 μg/ml pepstatin, 5 U/ml benzonase nuclease). Cell lysis was performed using a VS 70T sonotrode (20% amplitude; 0.7 sec pulse, 0.3 sec rest; 3× 30 sec with 1 min cooling in between) on a Sonoplus HD 3100 (Bandelin). Cell debris was removed by centrifugation at 50,000 × g for 30 min at 4°C. The supernatant was incubated with 2 ml of washed amylose beads for 15 min at 4°C and under constant rocking and then subjected to gravity-flow affinity columns. Unbound protein was washed away using 40 ml PP buffer. The first elution with 1 ml protein elution buffer (PP buffer containing 10 mM maltose) was discarded. The affinity purified protein was then eluted in 3 × 2 ml fractions. The protein concentration was determined by absorption spectroscopy using a NanoDrop 1000 spectrophotometer (PeqLab) at 280 nm with the protein specific molecular weight and the extinction coefficient calculated by the ExPASy ProtParam tool. The purified proteins were mixed with 5 × reducing protein sample buffer (8 M Urea, 0,15 % bromophenol blue, 5 mM EDTA, 10 % SDS, 0,1 M Tris-HCl, pH 6,8, 4 % Glycerol, 4 % β-mercapoethanol), boiled for 5 min at 95°C and subjected to SDS-PAGE in order to check the quality of the purified proteins. For further preparation and analytic purposes MBP-Opi1 variants were subjected to size exclusion chromatography using a Superdex 200 Increase 10/300 GL on an ÄKTA pure System (GE Healthcare). 500 μg protein were injected using a 500 μl loop and a flow rate of 0.5 ml/min in PP buffer. The void volume of the column (V_0_) was determined using BlueDextran eluting at 8.9 ml.

### Liposome flotation assays

Liposomes and protein variants were mixed at a molar protein:lipid ratio of 1:3300 in a total volume of 150 μl and incubated for 30 min at room temperature in a ultracentrifugation tube. For example, 80 μl of liposomes (4 mM) in NaPi buffer and 6 μg of protein in 70 μl PP buffer were mixed to yield 2.1 mM liposomes and 0.63 μM protein in 150 μl liposome flotation (LF) buffer (25 mM HEPES, 10 mM sodium phosphate, 150 mM sodium chloride and 0.5 mM EDTA, pH 7.4). After incubation, 100 μl of 75% sucrose dissolved in LF buffer were added and gently mixed with the sample to yield a final concentration of 30% sucrose. 200 μl of 20% sucrose dissolved in LF buffer were carefully layered on top of the 30% sucrose fraction and subsequently, 50 μl of LF buffer were layered on top of the 20% sucrose fraction, yielding in 500 μl of total volume. Sucrose density gradient centrifugation was conducted for 1 hour at 22°C at a speed of 240,000 × g in a Beckman TL-100 ultracentrifuge using a TLA 120.1 rotor (Beckman Coulter). Microlance™ 3 steel needles (Beckton, Dickinson) were used to collect four fractions of 125 μl from the bottom of the tube. Each fraction was mixed with 25 μl of 5 x reducing protein sample buffer, boiled for 5 min at 95°C and 10 μl of each fraction were subjected to SDS-PAGE using gradient gels (4.5-15 %, BioRad). Gels were stained with Instant Blue to visualize proteins as well as lipids. Protein amount of each fraction was quantified by densiometry using ImageJ. The bound fraction was determined as the amount of protein in the top fraction divided by the total protein content in all fractions together.

### Yeast viability assay and Opi^−^ test

Yeast strains were cultivated over night in 3 ml YPD (1% yeast extract, 2% peptone, 2% glucose) to an optical density (OD_600_) of ~10. 5 OD_600_ units of cells were washed twice with sterile water and spotted onto the respective plates using a metal stamp with a 1:10 dilution series starting with an OD_600_ of 1. Plates were scanned after 2 days of growth on solid synthetic complete dextrose (SCD) medium containing 75 μM inositol (+Ino) or lacking inositol (−Ino) at 30°C or 37°C.

For the Opi^−^ test, yeast strains were cultivated and washed with sterile water. 5 μl with an OD_600_ of 1 were spotted onto SCD plates lacking inositol. After 1 day of incubation at 30°C, an inositol auxotrophic tester strain (AID strain, kindly provided by Dr. Susan A. Henry, Cornell University, Ithaca, USA; washed and dissolved in sterile water) was sprayed on the plate followed by incubation for two additional days 30°C. Overproduction and secretion of inositol (Opi^−^ phenotype) was apparent from a red halo from the tester strain around the spotted yeast strains.

### Cell cultivation and calculation of growth rate

Yeast strains were cultivated over night in 3 ml SCD +Ino medium to reach an OD_600_ of ~7. A fresh culture was inoculated with the overnight culture to an OD_600_ of 0.33 in 3 ml SCD + Ino medium and cultivated for 4.5 hours. Cells were then harvested by centrifugation (4200 rpm, 3 min, 24°C), washed once with pre-warmed SCD -Ino and rediluted in 3 ml fresh and pre-warmed SCD -Ino medium for further cultivation at 30°C under constant agitation at 220 rpm. The OD_600_ was measured every 45 min up to 6.75 hours and the growth rate was calculated from the time window between three and six hours after the medium shift to guarantee inositol depleted conditions for all strains (Hofbauer *et al*, 2014; Gaspar *et al*, 2011).

### Fluorescence microscopy and image quantification

Cells were grown as described in *cell cultivation*. Microscopic images were recorded after 2 hours of inositol depletion on a Zeiss LSM770 confocal laser scanning microscope (Carl Zeiss AG) with spectral detection and a Plan-Apochromat 63 × 1.40 NA Oil DIC M27 objective. GFP fluorescence was excited at 488 nm and detected between 493-586 nm. RFP fluorescence was excited at 561 nm and detected between 578-696 nm. For better image visualization, the contrast was adjusted equally for all images using the ZEN 2 lite software (Carl Zeiss AG) with no further processing.

For quantification, the raw images were smoothed with a Gaussian filter of radius 2. To extract the nucleus-related structures, either nuclear localization or nuclear ER localization, the objects with highest intensities were identified by an intensity clustering algorithm. All objects above a size threshold were classified as nucleus-related structures. The size threshold was manually adjusted for the different conditions and ranged between 0.73 μm² and 2.15 μm². For each nucleus-related structure, the localization index 
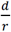
 was calculated, where *d* is the maximum distance of the pixels of an object to the centroid of the object and *r* is the equivalent disk radius, here particularly the radius of a disk consisting of the same number of pixels as the object of interest. If the fluorescent particles have a nuclear localization, the observed shape is a blob with the size of the nucleus. The equivalent disk radius *r* of a blob is comparable to *d* and the localization index is close to one. If the fluorescent particles localize to the ER, one observes a ring-shaped structure around the nucleus. For a ring, the maximum distance to the centroid is bigger than the radius of an area-equivalent disk (*r* < *d*) and the localization index is large. We have set the threshold between blob and ring to 1.3. Hence, a localization index of up to 1.3 indicates nuclear localization. For a localization index above 1.3, we have localization to the nuclear ER. For each condition, we analyzed at least 190 cells in 4 images. The image analysis was implemented in Mathematica 11.1 (Wolfram Research). The code and an example calculation are available at https://www.physikalischebiologie.de/downloads.

### Analysis of Opi1-mGFP protein levels using immunoblotting

Cells were grown as described in *cell cultivation*. 2 hours after the shift to medium lacking inositol, 3 OD_600_ units were harvested (3500 rpm, 5 min, 4°C) and washed once with ice-cold water. Cell pellets were resuspended in 400 μl 12.5% trichloroacetic acid (TCA) and stored at –20°C over night. TCA was removed by centrifugation at 13,000 rpm, 5 min, room temperature, and the cell pellets were washed with 500 μl 80% ice-cold acetone, air-dried for 2 min, and dissolved in 160 μl of a mixture of 1% SDS and 0.1 N NaOH. 40 μl of 5 × protein sample buffer were added, samples were boiled at 95°C for 5 min. SDS-PAGE was performed using 7.5% gels (BioRad) and proteins were blotted onto nitrocellulose membranes (Roth) and incubated with 5% skim milk powder (Fluka) in TBST (50 mM Tris, 150 mM sodium chloride, pH 8.0, 0.1% Tween-20) for 30 min to reduce unspecific antibody binding. Blots were then decorated with 2.5% skim milk powder in TBST containing either mouse anti-GFP antibody (11-814-460-001, Roche Diagnostics; 1:1000 dilution) or mouse anti-Pgk1 antibody (22C5D8, Life Technologies; 1:20000 dilution) as primary antibodies and anti-mouse-HRP antibody (115-035-003, Dianova; 1:20000 dilution) as secondary antibody for detection with SuperSignal^®^ West Femto Maximum Sensitivity Substrate (Thermo Scientific) using the ChemiDoc^TM^ MP Imaging System (BioRad).

### Coarse-grained molecular dynamics simulations

We modelled a 22-mer peptide (QKLSRAIAKGKDNLKEYKLNMS) into an α-helical conformation using the UCSF CHIMERA software package (Pettersen *et al*, 2004). In a coarse-grained (CG) representation using in the MARTINI force field (Marrink *et al*, 2007; De Jong et al, 2013), the helix was placed in a cubic box of 10x10x10nm^3^ containing water, a lipid bilayer and 150 mM sodium chloride, assembled using the insane tool (Wassenaar *et al*, 2015). The helix was placed at a distance of 4 nm from the bilayer’s center of mass. We set up different bilayer systems with different lipid compositions, all spanning the xy-plane. The bilayer system designated as 20% PA was composed of 60 mol% POPC, 20 mol% DOPC, and 20 mol% DOPA. The bilayer designated as 20% PS was composed of 60 mol% POPC, 20 mol% DOPC, and 20 mol% DOPS. The bilayer designated as 20% PA/PS contained 60 mol% POPC, 20 mol% DOPA, and 20 mol% DOPS. Acyl chains were kept constant with a ratio of 6:4 (PO/DO), while the head group species and their relative ratios were varied. Each system was simulated for 1 μs.

CG simulations were performed using the GROMACS software package version 5.1.3 (Abraham *et al*, 2015) with the MARTINI force field version 2.2. The integration time step was set to 20 fs with a neighbor list update every 20 steps. Non-bonded interactions were cut off at 1.1 nm using the Verlet algorithm. Temperature was kept constant at 310 K using the Velocity Rescale (Bussi *et al*, 2007) thermostat with a characteristic time of 1 ps. A pressure of 1 bar was maintained by semiisotropic pressure coupling using the Parrinello-Rahman algorithm and a characteristic time of 12 ps (Parrinello & Rahman, 1980). Periodic boundary conditions were imposed.

### Backmapping of coarse-grained systems to an atomistic representation

We used CHARMM-GUI (Jo *et al*, 2007, 2008, 2009, 2014; Lee *et al*, 2016; Wassenaar *et al*, 2014) to lift all three CG systems to an atomistic description in the CHARMM36 force field (Best *et al*, 2014; Klauda *et al*, 2010; Vanommeslaeghe *et al*, 2010). 150 mM NaCl and neutralizing counterions were added. The energy of the system was minimized for 5000 steps, keeping a restraint on all heavy atoms. NVT-ensemble equilibration was run for 50 ns at a temperature at 310 K using a time step of 1 fs. Subsequently an NPT-ensemble equilibration was run for 375 ns with a time step of 2 fs to stabilize the pressure at 1 bar. Both the Berendsen barostat and thermostat were used during equilibration (Berendsen *et al*, 1984). Non-bonded interactions were cut off at 1.2 nm. During the production simulations we used Velocity Rescale thermostat (Bussi *et al*, 2007) at 310 K and the Parrinello-Rahman barostat (Parrinello & Rahman, 1980) with semi-isotropic pressure coupling, applying them on the protein, membrane, and solvent with characteristic times of 1 and 5 ps, respectively. We simulated the systems for 5 μs each, using the last 4.9 μs for data analysis.

### Analysis of lipid localization

For every frame in the trajectory we binned phosphate atoms belonging to a specific lipid type according to their x and y positions on a two-dimensional grid, using a bin area of 1 Å^2^ and sampling every 500 ps. We then obtained a 2D in-plane rotation angle for every frame by using an in-plane least-square fit of the backbone C_α_ atoms to a reference structure of the amphipathic helix (AH) aligned along the x-axis. For every frame, the two-dimensional grid was interpolated by a third order spline and rotated by the calculated angle about the direction orthogonal to the plane of the membrane. By considering periodic boundary conditions, the grid in every frame was extended and cropped after the rotation to keep the original dimensions. Finally, we calculated the lipid density by averaging all two-dimensional grids along the last 4.9 μs.

### Calculation of distance distributions

We centered the AH at the origin and aligned the AH on the x-axis with the N-terminus pointing in the positive x direction. We then calculated the pairwise distances from a reference point to all lipids localized in a given area. We defined the combined Cα center of mass of three amino acids (K112, R115 and K119 for the KRK motif and K121, K125 and K128 for the 3K motif) as our reference points p_KRK_ and p_3K_. We then defined two boundaries px ± 10Å on the x-axis from each point of reference. For p_3K_ we considered lipids between the boundaries with negative y, while for p_KRK_ we considered lipids with positive y values. We then calculated for every frame the 2D Euclidean distance to all lipids contained in the squared area adjacent to the points of interest. We binned all calculated distances into 200 bins, using a bin width of 0.1 Å, sampling every 500 ps.

### Residence times

We calculated for each trajectory the residence time of lipids in the neighborhood of residues K112, R115, and K119, or K121, K125, and K128. We removed the coordinates of the center of mass of all atoms and aligned the AH onto the x-axis as described above. We defined three rectangular areas A_b_, A_i_ and A_u_ around the region of interest p and in each frame assigned a state S to each lipid based on its localization in a given area: S_b_ (bound), S_i_ (intermediate) or S_u_ (unbound). We considered a lipid to be bound when first entering region Ab and to dissociate when first hitting region Au. We also defined an intermediate region Ai in which the lipids commit for association or disassociation. We calculated the residence time by summation of the total time spent in state S_b_ and S_i_ before hitting state S_u_ again (Buchete & Hummer, 2008).

### Trajectory analysis

We used VMD (Humphrey *et al*, 1996), GROMACS (Abraham *et al*, 2015), MDAnalysis (Michaud-Agrawal *et al*, 2011; Gowers *et al*, 2016), NumPy (Van Der Walt *et al*, 2011), SciPy (Jones *et al*, 2001), IPython (Pérez & Granger, 2007) and Matplotlib (Hunter, 2007) for the analysis and visualization of trajectories.

---

**Expanded view Figure EV1.**
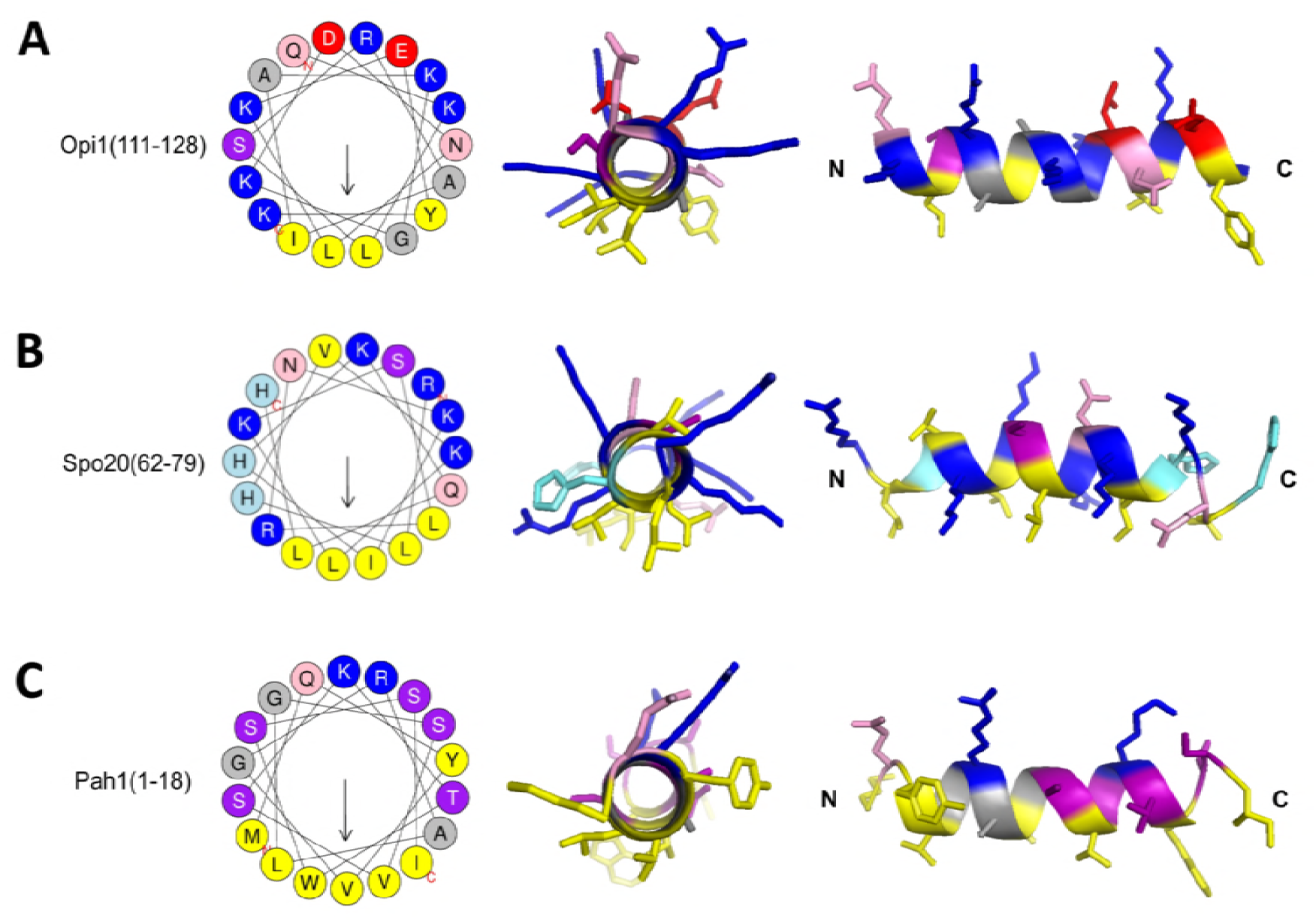
Amphipathic helices of proteins sensing PA-rich membranes. A-C Visualization of the AH of Opi1 (A), Spo20 (B) and Pah1 (C) using the Heliquest tool (Gautier et al., 2008) and the PyMOL^TM^ Molecular Graphics System for front and side views of the helix matching the color code of the Heliquest tool.

**Expanded view Figure EV2.**
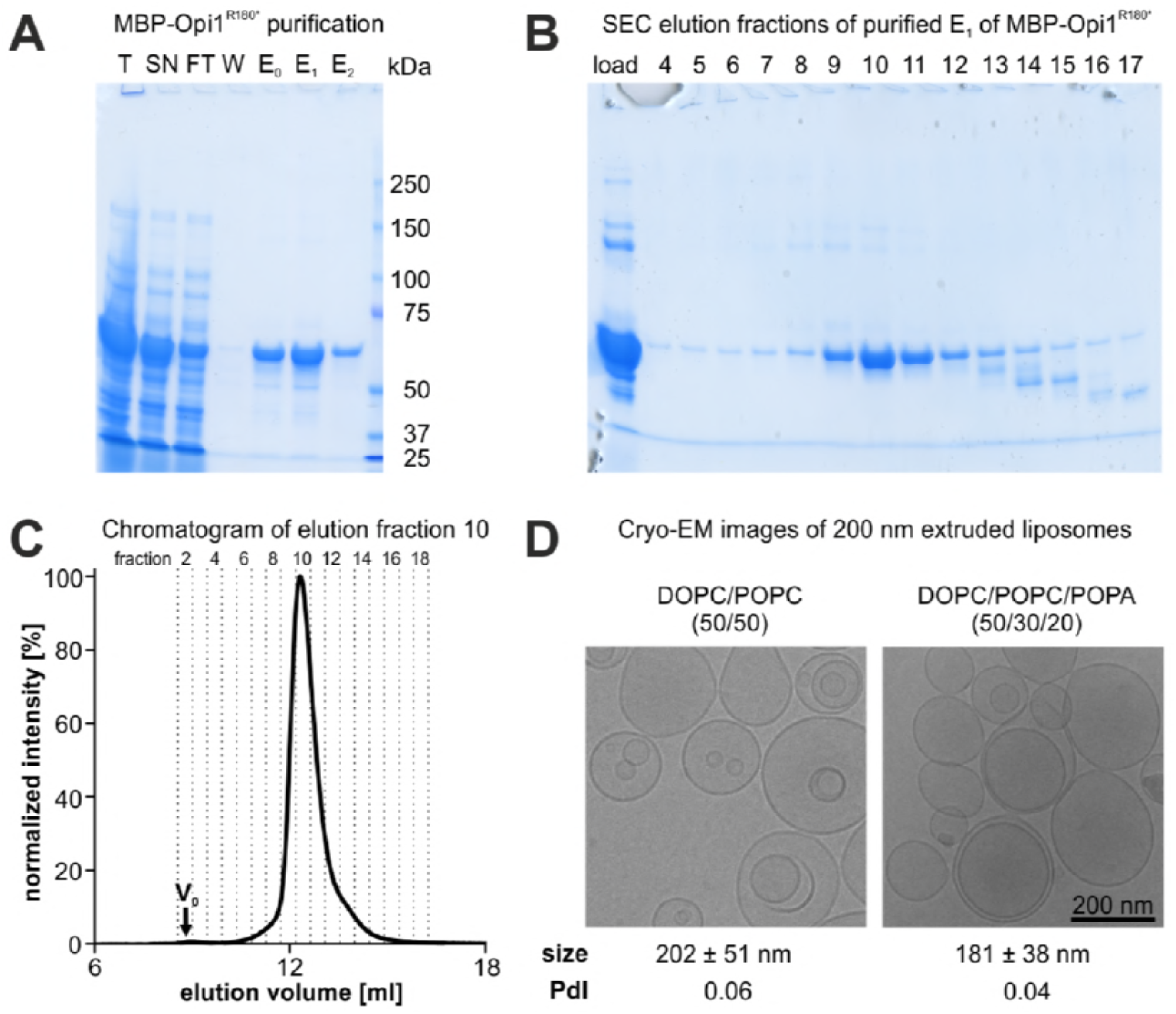
Quality controls of purified MBP-Opi1R^180*^ variants and liposomes. A Purification of MBP-Opi1R^180*^; SDS-PAGE of sample fractions from the affinity purification. B SDS-PAGE of size exclusion chromatography (SEC) elution fractions from the affinity purified elution fraction E_1_ of MBP-Opi1R^180*^. C Quality control of SEC purified protein; SEC elution fraction 10 of MBP-Opi1R^180*^ was again subjected to size exclusion chromatography to verify good and stable protein quality. The void volume (V_0_) of the Superdex 200 Increase 10/300 GL column is 8.9 ml. D Cryo-electron microscopic images of liposomes generated by extrusion through a 200 nm pore size polycarbonate filter. Scale bar, 200 nm. NanoSight measurements were performed to determine the mean size +/– SD and the polydispersity index (PdI) of the extruded liposomes.

**Expanded view Figure EV3.**
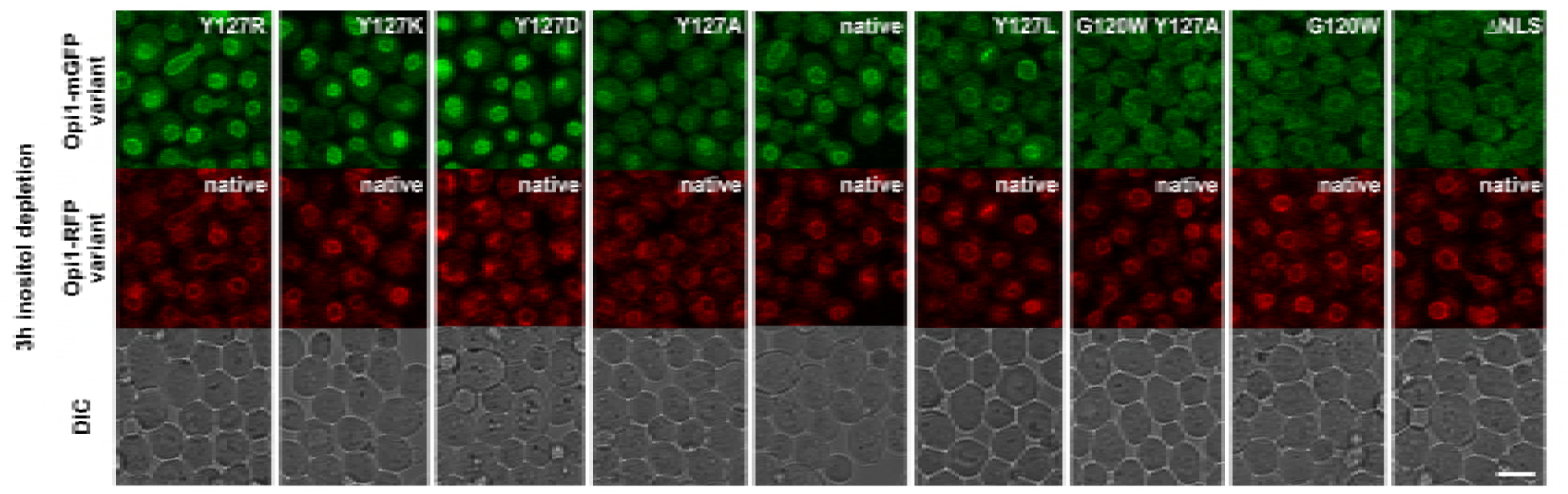
Opi1 point mutations do not impair nuclear ER membrane binding of native Opi1 in diploid yeast cells. Representative microscopic images of diploid strains with chromosomally integrated Opi1-mGFP mutant variants as one allele and wild type Opi1-mRFP as the other allele. The cells were cultivated and imaged three hours after inositol depletion in liquid media. Scale bar, 5 μm.

**Expanded view Figure EV4.**
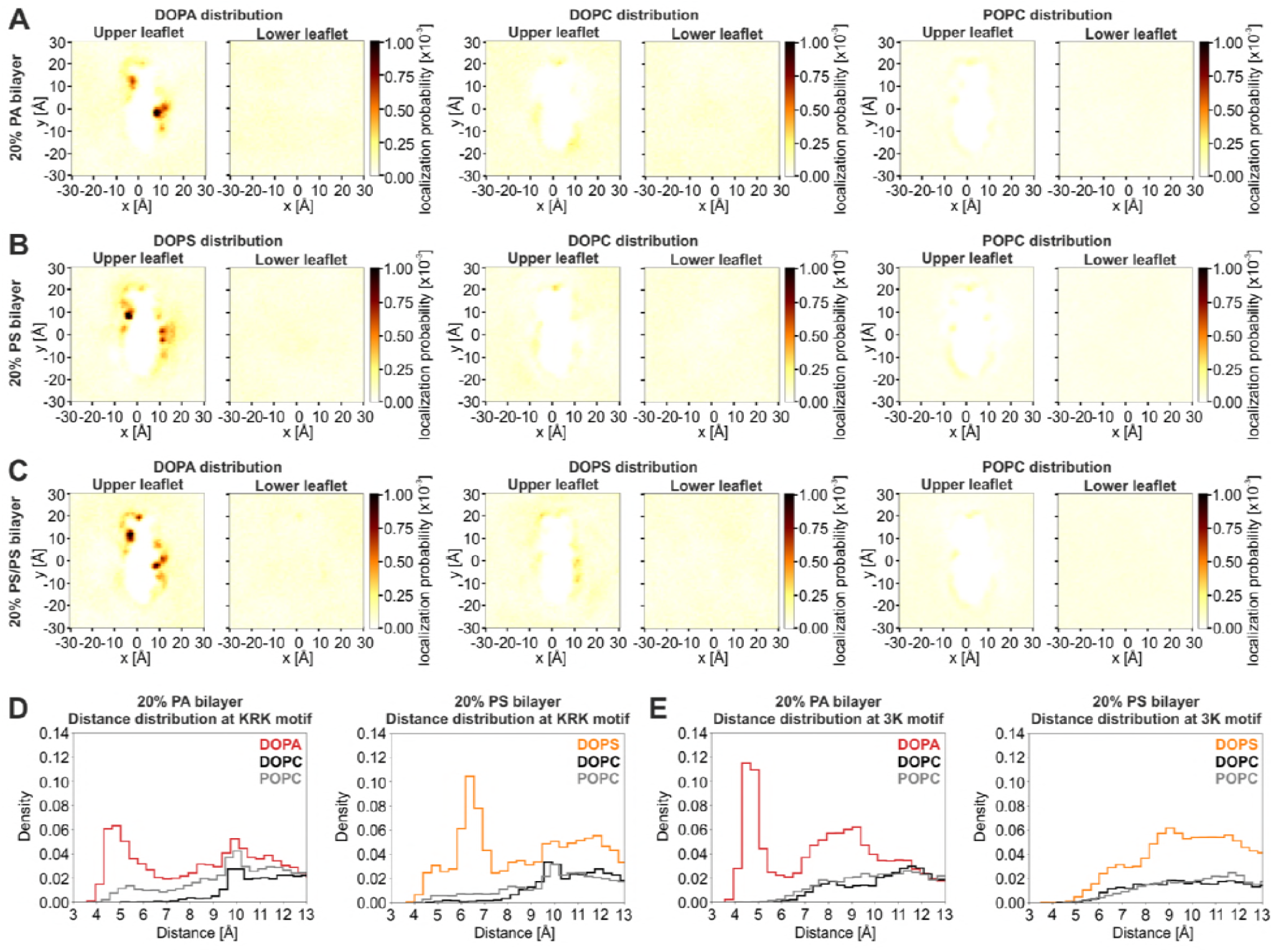
A-C Time-averaged positions of phosphate head groups from different lipid species (DOPA, DOPS, DOPC, and POPC) in their respective all-atom MD simulations (A: 20% PA, B: 20% PS, C: mixed 20% PA / 20% PS). Both membrane leaflets are shown: upper leaflet (Opi1 AH bound) and lower leaflet (unbound). Colors indicate the probability to observe a given lipid at that position over the course of the trajectory. DOPA and DOPS show enrichment at the 3K- and KRK-motifs in their respective simulations (A and B, left panels), whereas no enrichment is seen for DOPC and POPC (A and B, middle and right panels). DOPA displaces DOPS from both motifs in the mixed simulation (C, left and middle panels). D,E Distribution of pairwise distances calculated from a point of interest and all lipids of a given species in their respective simulations. (D) Distribution of lipids at the KRK-motif for DOPA, DOPC, and POPC (left panel) and DOPS, DOPC and POPC (right panel). (E) Distribution of lipids at the 3K-motif for DOPA, DOPC, and POPC (left panel) and DOPS, DOPC, and POPC (right panel). The point of interest was defined as described in Figure 5G/5I.

**Table EV1.**
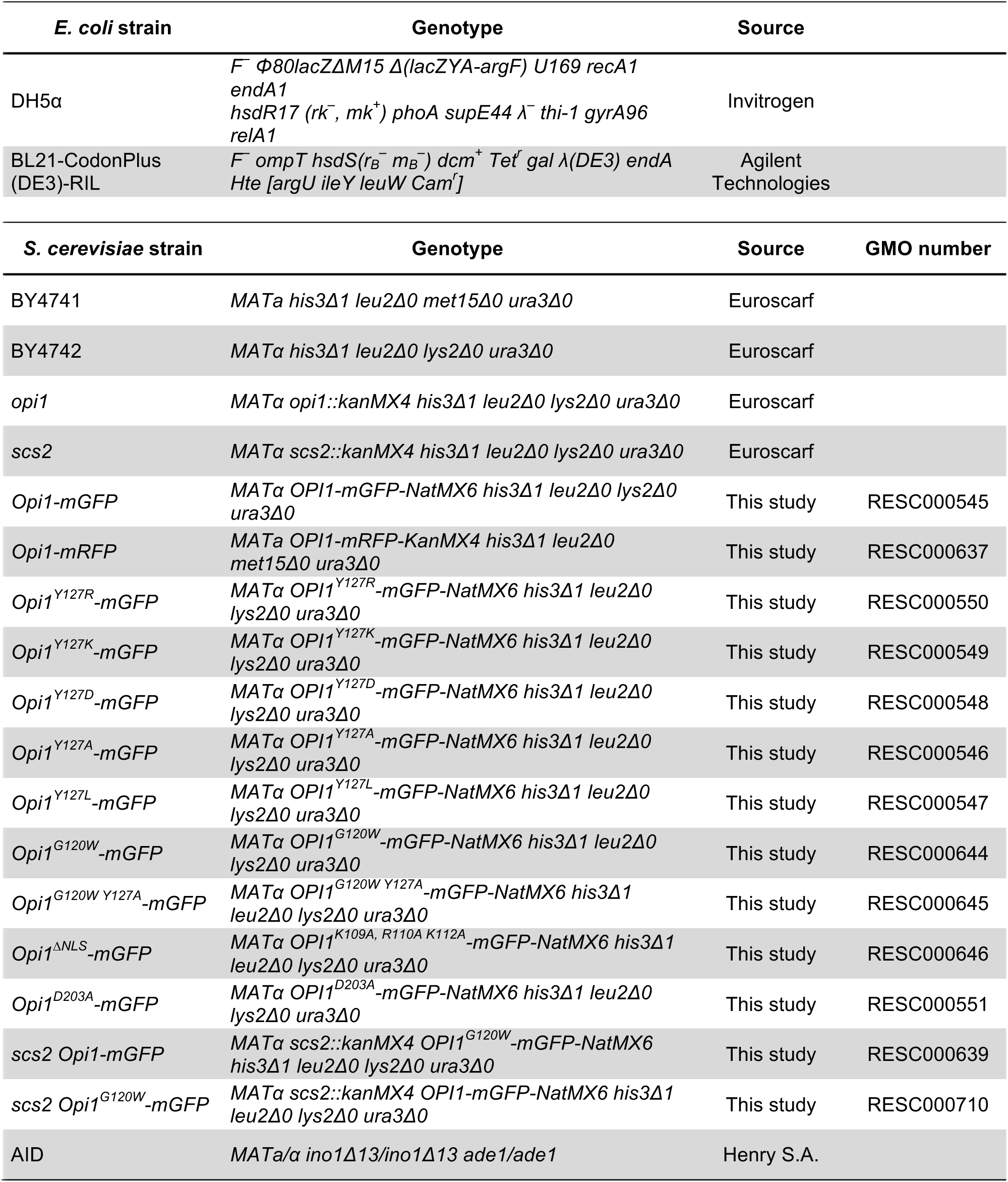
*E. coli* and *Saccharomyces cerevisiae* strains used in this study.

**Table EV2.** Plasmids used in this study.

| Name | Bacterial marker | Source | Plasmid number |
| --- | --- | --- | --- |
| pMAL-Opi1 | Amp | Carman G.M. | pRE394 |
| pFA6a-mGFP-NatMX6 | Amp | Kohlwein S.D. | pRE390 |
| pFA6a-mRFP-KanMX4 | Amp | Kohlwein S.D. | pRE391 |
| pMAL-Opi1<sup>S114*</sup> | Amp | This study | pRE634 |
| pMAL-Opi1<sup>R180*</sup> | Amp | This study | pRE636 |
| pMAL-Opi1<sup>Y127A,R180*</sup> | Amp | This study | pRE660 |
| pMAL-Opi1<sup>Y127D,R180*</sup> | Amp | This study | pRE661 |
| pMAL-Opi1<sup>Y127W,R180*</sup> | Amp | This study | pRE662 |
| pMAL-Opi1<sup>G120W,R180*</sup> | Amp | This study | pRE663 |
| pMAL-Opi1<sup>2xAH,R180*</sup> | Amp | This study | pRE664 |
| pMAL-Opi1<sup>D122K,E126K,R180*</sup> | Amp | This study | pRE665 |
| pMAL-Opi1<sup>5K→5R,R180*</sup> | Amp | This study | pRE676 |
| pFA6a-Opi1-mGFP | Amp | This study | pRE397 |
| pFA6a-Opi1<sup>Y127A</sup>-mGFP | Amp | This study | pRE587 |
| pFA6a-Opi1<sup>Y127L</sup>-mGFP | Amp | This study | pRE398 |
| pFA6a-Opi1<sup>Y127D</sup>-mGFP | Amp | This study | pRE399 |
| pFA6a-Opi1<sup>Y127R</sup>-mGFP | Amp | This study | pRE589 |
| pFA6a-Opi1<sup>Y127K</sup>-mGFP | Amp | This study | pRE588 |
| pFA6a-Opi1<sup>G120W</sup>-mGFP | Amp | This study | pRE591 |
| pFA6a-Opi1<sup>G120W,Y127A</sup>-mGFP | Amp | This study | pRE592 |
| pFA6a-Opi1<sup>K109A,R110A,K112A</sup>-mGFP | Amp | This study | pRE593 |
| pFA6a-Opi1<sup>D203A</sup>-mGFP | Amp | This study | pRE590 |

**Table EV3.**
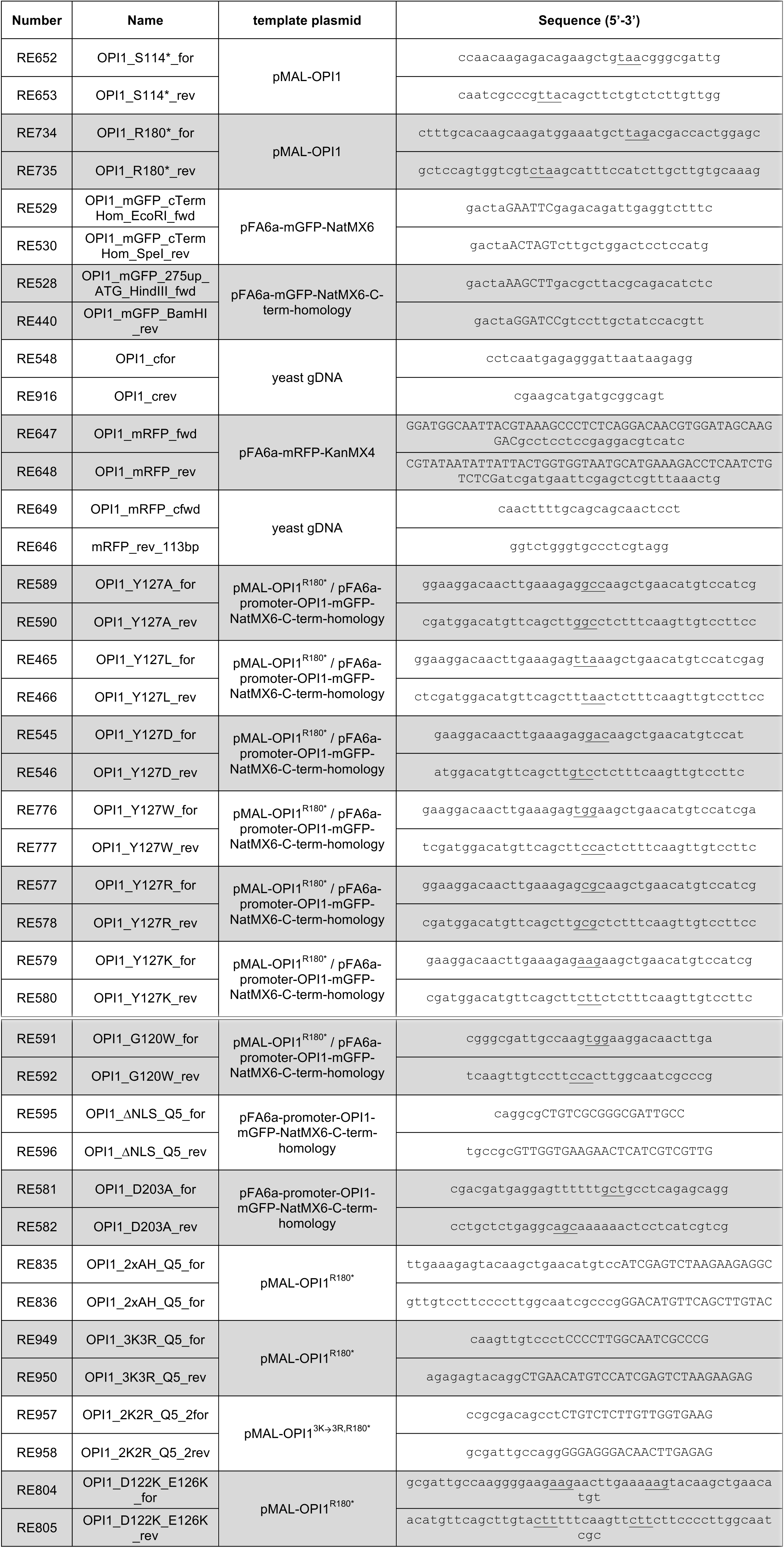
Primers used in this study.

